# Cholinergic–dopaminergic synergy and antagonism shape prediction-error broadcasting

**DOI:** 10.64898/2026.02.19.706866

**Authors:** Bálint Király, Vivien Pillár, Írisz Szabó, Dániel Schlingloff, Panna Hegedüs, Krisztián Szigeti, Yulong Li, Balázs Hangya

## Abstract

Cholinergic and dopaminergic pathways control reinforcement learning, yet how their basal forebrain and midbrain populations interact remains unresolved. We simultaneously monitored both systems using dual-cell-type optogenetic tagging and neurotransmitter measurements in mice engaged in psychometric learning. Their activity contributed synergistically to reward prediction-error signaling. However, punishment-inhibited dopaminergic neurons showed antagonistic interactions with cholinergic neurons as well. Optogenetic stimulation and channelrhodopsin-assisted circuit mapping revealed a cholinergic-to-dopaminergic disynaptic inhibitory pathway through basal forebrain GABAergic neurons consistent with this antagonism. Chemogenetic suppression of cholinergic activity reduced dopaminergic responses to reward-predicting stimuli, attenuated punishment-induced dopamine suppression, and impaired learning. These findings reveal how a common underlying circuit architecture can flexibly shape prediction-error signaling through synergistic and antagonistic interactions, with implications for adaptive learning and neuropsychiatric disease.

## Introduction

Neuromodulatory systems play a central role in reinforcement learning, with the dopaminergic system encoding reward prediction error (RPE) – the difference between experienced and expected reward (*1*). This notion has profoundly shaped the study of neuromodulators, including acetylcholine (ACh) and noradrenaline, which have often been assigned distinct computational roles such as learning rate and inverse temperature, respectively (*2*, *3*).

However, recent studies have shown that the central cholinergic system is also involved in encoding outcome expectations and can be described as carrying an unsigned prediction error (UPE) signal – the absolute value of RPE (*4*, *5*). These findings raise the question of how cholinergic and dopaminergic populations coordinate to shape prediction-error signaling during learning. Do cholinergic and dopaminergic populations cooperate or compete, and are potential synergistic or antagonistic interactions fixed or dynamically organized across behavioral epochs?

In computational terms, this question can be addressed by examining how cholinergic and dopaminergic activity covaries beyond their average event-related responses and how their interactions unfold across temporal scales (*6*). Noise correlations capture moment-by-moment covariation between the two systems, with positive correlations consistent with synergistic interactions and negative correlations indicating antagonistic interactions. More broadly, synergistic interactions may emerge when the two systems jointly contribute information beyond that carried by either system alone. Critically, resolving these interactions requires simultaneous recordings of cholinergic and dopaminergic activity in behaving animals with sufficient temporal precision to capture both within-trial interactions and learning-related dynamics across trials. We therefore performed dual-cell-type optogenetic tagging of basal forebrain cholinergic neurons (BFCNs) and midbrain dopaminergic neurons (DANs), and fiber photometry recordings of neurotransmitter release. We found that cholinergic activity engaged distinct dopaminergic subtypes through temporally organized synergistic and antagonistic interactions, thereby uncovering a cell-type-specific interaction architecture that was underpinned by a disynaptic cholinergic-to-dopaminergic circuit.

## Results

### Shared prediction-error signaling across cholinergic and dopaminergic populations

To investigate how cholinergic and dopaminergic populations coordinate during reinforcement learning, we trained mice on a head-fixed auditory operant conditioning task featuring both stable and changing associations (Fig. 1A). Mice first learned to lick in response to a tone paired with water reward and to withhold licking to another tone paired with air-puff punishment; these associations were kept fixed throughout the training. We then introduced novel tone-outcome associations in randomized order. Some novel associations resembled the original contingencies and were learned readily, whereas others differed substantially and were learned more slowly, generating a psychometric relationship between learning speed and similarity to the original associations (Fig. 1B). We refer to this behavioral paradigm as the psychometric learning task. Throughout training, mice acquired the novel associations while maintaining stable performance on the fixed contingencies (Fig. S1; 1-6 novel pairings introduced per session, mean 2.29). The coexistence of stable and newly acquired associations allowed us to dissociate responses reflecting well-established expectations from those accompanying the formation of new predictions, providing a behavioral context in which cholinergic–dopaminergic interactions could be examined both moment-to-moment and across learning.

**Figure 1.**
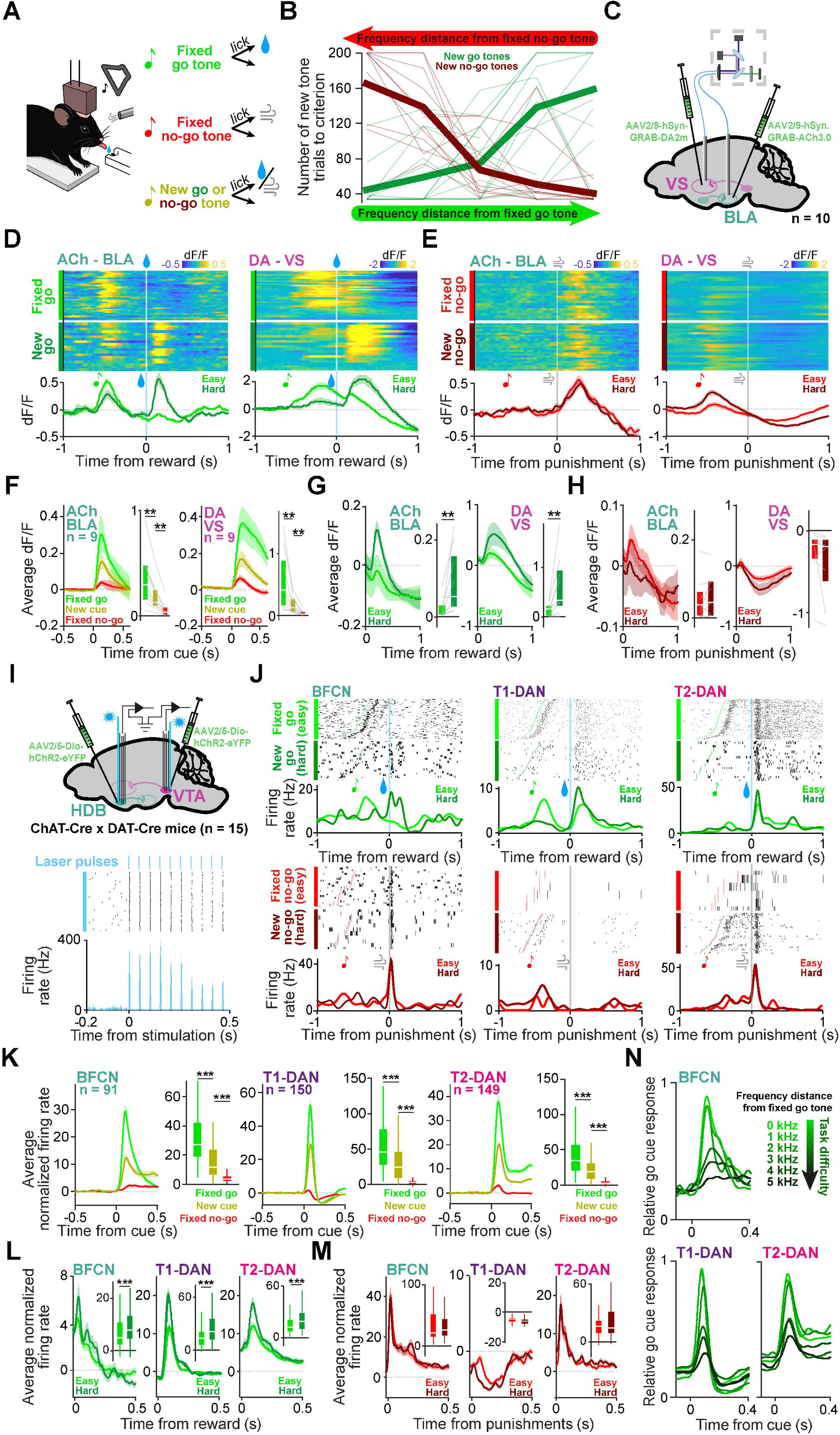
Dopamine and acetylcholine represent outcome prediction error signals. **A** Schematic of the auditory operant go/no-go task. Head-fixed mice learned to lick for water reward after go tones and withhold licking to avoid air puff punishments after no-go tones. Fixed go and no-go tones were complemented by novel tone-outcome pairings that were randomly updated every time mice reached 70% performance after a minimum of 33 trials between switches. **B** Psychometric curves show that learning speed, indexed by the average number of trials required to reach criterion performance (capped at 200, when the criterion was not reached), was correlated with difficulty, indexed by the frequency distance of the new tone from the fixed tone predicting the same outcome (thin lines, individual animals; thick lines, grand averages). **C** Schematic of simultaneous fiber photometry recordings of ACh release in the BLA and DA release in the VS. **D** Normalized fluorescent signals (dF/F) of the ACh sensor in the BLA (left) and DA sensor in the VS (right), aligned to reward delivery in an example session. Top, single trials grouped by task difficulty (‘easy’, rewards following fixed tones; ‘hard’, rewards following novel tones). Bottom, session average. **E** Same as D, but aligned to punishment delivery. **F** Grand average fluorescent signals of ACh sensors in the BLA (left) and DA sensors in the VS (right) aligned to cue tone presentations, partitioned by cue tone types (n = 9 mice). Left, peri-event time histograms (PETHs); right, mean response magnitudes. **G-H** As in panel F, but aligned to rewards (G) and punishments (H). ACh release was consistent with an unsigned, while DA release with a signed RPE representation. **, p = 0.0039; two-sided Wilcoxon signed-rank tests. **I** Schematic of concurrent electrophysiological recordings of HDB cholinergic and VTA dopaminergic neurons. Bottom, spike raster and PETH of an optogenetically identified dopaminergic neuron, aligned to blue laser pulses (1ms pulses @20Hz). **J** Top, example spike rasters of BFCN (left), T1-DAN (middle) and T2-DAN (right) neurons aligned to reward (blue line, top) and punishment (gray line, bottom) delivery. Single trials were grouped by task difficulty. Color ticks indicate cue tone onset times. Bottom, PETHs of the same example recordings smoothed with a Gaussian kernel (width, 20 ms). **K** Grand average PETH of BFCN (left, n = 91 neurons), T1-DAN (middle, n = 150 neurons) and T2-DAN firing (right, n = 149 neurons) aligned to cue tone presentations, partitioned by cue tone types. Left, peri-event time histograms (PETHs); right, mean response magnitudes. **L-M** As in panel K, but aligned to reward (L) and punishment (M) delivery, partitioned by task difficulty. BFCN activity was consistent with an unsigned, and T1-DAN activity with a signed RPE representation, congruent with the neuromodulator release shown in panels F-H; however, T2-DAN activity showed the hallmarks of unsigned prediction errors, similar to BFCNs and unlike T1-DANs. ***, p < 0.001; two-sided Wilcoxon signed-rank tests (BFCN: fixed go vs. novel, p = 1.2 x 10^-16^; novel vs. fixed no-go, p = 5.6 x 10^-15^; easy reward vs. hard reward, p = 6.7 x 10^-5^. T1-DAN: fixed go vs. novel, p = 3.4 x 10^-26^; novel vs. fixed no-go, p = 3.3 x 10^-25^; easy reward vs. hard reward, p = 7.9 x 10^-14^. T2-DAN: fixed go vs. novel, p = 3.4 x 10^-26^; novel vs. fixed no-go, p = 1.3 x 10^-25^; easy reward vs. hard reward, p = 2.9 x 10^-13^). **N** Average normalized PETHs aligned to go cues. BFCN, T1-DAN and T2-DAN responses scaled with task difficulty, indexed by the frequency distance of the cue from the fixed go tone. Box-whisker plots show median, interquartile range and non-outlier range. Error shades indicate SEM.

To compare subcortical cholinergic and dopaminergic prediction-error signalling during learning, we simultaneously expressed the acetylcholine-sensor GRAB-ACh3.0 in the basolateral amygdala (BLA), a major target of BFCNs receiving prediction error signals (*4*, *7*, *8*), and the dopamine-sensor GRAB-DA2m in the ventral striatum (VS), the primary locus of dopaminergic RPE signaling (Fig. 1C, S2B and S2D) (*9–11*). Reward-predicting stimuli evoked phasic release of both ACh and DA, scaled by the reward-predicting value of the cue (fixed go > new cues > fixed no-go; Fig. 1D-F and S3). Reward delivery also triggered release of both transmitters, with less-expected rewards on ‘hard’ trials during new learning eliciting larger responses than confidently expected rewards associated with fixed stimuli, consistent with RPE coding (*10*) (Fig. 1D,G). A key difference emerged for punishments: they increased ACh release but suppressed DA release (Fig. 1E,H), confirming that cholinergic prediction error signals are unsigned (*4*, *12–15*). Thus, cholinergic and dopaminergic release exhibited similar sensitivity to reward-predicting cues and rewards while diverging in their responses to punishment.

To examine cholinergic-dopaminergic interactions with single-neuron and millisecond temporal resolution, we simultaneously recorded cholinergic and dopaminergic neurons during learning. We generated a double transgenic ChAT-Cre × DAT-Cre mouse line expressing Cre recombinase in both cholinergic and dopaminergic neurons (Fig. S4A). We expressed channelrhodopsin-2 by viral gene transfer in the horizontal limb of the diagonal band (HDB) and the ventral tegmental area (VTA), allowing optogenetic identification of both neuronal populations during extracellular recordings while mice performed the psychometric learning task (Fig. 1I, S2A,C,E and S4B–D) (*12*, *16–18*). Because neither direct HDB-to-VTA cholinergic nor VTA-to-HDB dopaminergic projections exist (Fig. S4E,F, (*19–21*)), fast indirect activation of one population by photostimulation of the other could not confound optogenetic identification. Consistent with this, we detected no cross-activation (Fig. S4B). Clustering of neuronal response properties similar to earlier studies (*22–24*) identified putative cholinergic (n = 91) and dopaminergic (n = 299) populations, yielding n = 64 simultaneously recorded cholinergic-dopaminergic pairs for subsequent analyses (Fig. S5).

Cholinergic and dopaminergic neurons exhibited structured responses to reward-predicting cues and outcomes, consistent with prediction-error signaling (Fig. 1J-M). BFCNs responded strongly to fixed go cues, followed by novel cues and fixed no-go cues, reflecting the reward-predictive value of the auditory stimuli. They were activated by both rewards and punishments, consistent with UPE coding. Many DANs (Type 1, T1-DANs, n = 150) showed similar gradation of cue-evoked activity, activation after rewards, but inhibition after punishments, consistent with RPE coding. By contrast, about half the dopaminergic neurons (Type 2, T2-DANs, n = 149) were activated after punishments (*10*, *24–27*), yielding response profiles that qualitatively resembled those of BFCNs. Although the two dopaminergic types were intermingled in the VTA, T2-DANs were preferentially located in the medial and ventral VTA and near the substantia nigra pars compacta (SNc; Fig. S2E), consistent with earlier studies (*28–31*).

We next asked whether cholinergic and dopaminergic populations shared prediction-error coding across multiple dimensions of learning. In both BFCNs and DANs, cue responses scaled with task difficulty (Fig. 1N and S6A), whereas reward responses scaled inversely with reward expectation. Confidently expected ‘easy’ rewards elicited smaller trial-averaged responses than less-expected rewards on ‘hard’ trials (Fig. 1L). Punishments, which are largely unexpected in operant tasks (Fig. 6 in ref. (*12*)), evoked relatively uniform responses (Fig. 1M). Response magnitude also tracked trial-by-trial behavioral confidence: responses to rewards were smaller when animals responded faster (Fig. S6B). Consistent with temporal-difference models, which predict larger prediction errors for outcomes occurring earlier or later than expected (*32*, *33*), both neuromodulatory systems responded more strongly when feedback deviated from the average delay (100–300 ms; Fig. S6C,D). Finally, correct rejections were followed by activation of both neuromodulatory systems, extending prediction-error signaling to successful avoidance of aversive events (Fig. S7) (*34*, *35*). Together, these results show that cholinergic and dopaminergic populations tracked prediction-error signals across multiple dimensions of outcome expectation, raising the question of how their shared prediction-error signaling is coordinated through interactions between the two systems.

### Cell-type- and task-phase-dependent segregation of cholinergic-dopaminergic interactions

We next asked how cholinergic and dopaminergic populations coordinate beyond their trial-averaged responses, focusing on moment-by-moment covariation in neuronal activity. We first examined these interactions using an unbiased spike-time cross-correlation analysis that included all recorded spikes. This analysis revealed distinct interaction signatures for the two dopaminergic populations: T1-DANs exhibited a short-latency suppression following cholinergic spikes, whereas T2-DANs showed a zero-lag positive correlation with cholinergic firing (Fig. 2A). The asymmetric cholinergic suppression of T1-DANs was also evident outside the task context, indicating that this interaction extends beyond task-evoked activity (Fig. S8A).

**Figure 2.**
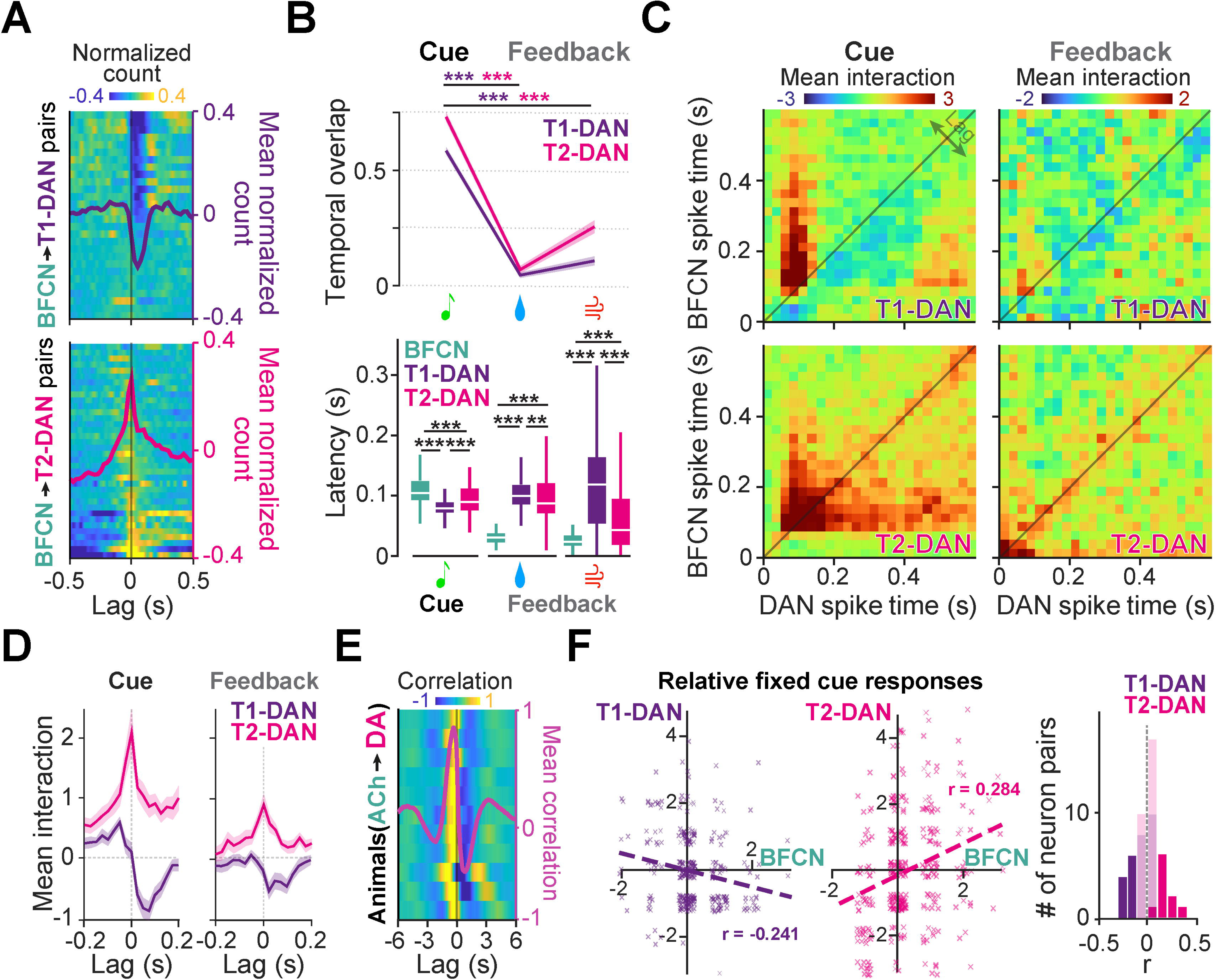
Segregation of cholinergic–dopaminergic interactions by dopaminergic cell type and behavioral context. **A** Top, cross-correlograms of BFCNs and T1-DANs (rows, pairs of concurrently recorded neurons; blue, low correlation; yellow, high correlation) showing short-latency suppression of T1-DAN activity after BFCN spikes. Bottom, cross-correlograms of BFCNs and T2-DANs showing zero-lag co-activity. Averages are overlaid. **B** Top, mean Jaccard index of event-evoked BFCN and DAN response intervals. Error shades, SEM. Bottom, distributions of peak response latencies. Cue responses showed larger temporal overlap and a T1-DAN → T2-DAN → BFCN sequence, whereas feedback responses showed smaller overlap and the reverse sequence (two-sided Wilcoxon signed-rank tests; **, p < 0.01; ***, p < 0.001). Box-whisker plots show median, interquartile range and non-outlier range. **C** Time-resolved BFCN–DAN interaction matrices (JPETH minus the product of the PETHs, see Fig. S8E,F; blue, negative; red, positive) averaged over all simultaneously recorded BFCN – T1-DAN (top) and BFCN – T2-DAN pairs (bottom) aligned to cue (left) and feedback (right). **D** Mean BFCN–DAN interaction as a function of time lag between neurons for cue-evoked (left) and feedback-evoked (right) activity. Error shades indicate SEM. **E** Cross-correlogram of ACh and DA release in the BLA and the VS, respectively (rows, session averages per mouse; blue, negative correlation; yellow, positive correlation). ACh release correlated best with subsequent release of DA (−0.36 ± 0.31 s mean peak lead ± standard deviation across animals; significantly non-zero, p = 0.0039 two-sided Wilcoxon signed-rank test). The curve shows the grand average across animals. **F** Left, trial-by-trial correlation of cholinergic vs dopaminergic responses to fixed cues relative to the neuron’s average, for example pairs of neurons. Relative responses were significantly correlated (Spearman correlation, BFCN—T1-DAN example, r= −0.241, p = 3.9 x 10^-5^; BFCN—T2-DAN example, r = 0.284, p = 4.0 x 10^-8^). Dashed lines show fitted linear regression. Right, histogram of the Spearman correlation coefficients of all concurrently recorded cholinergic-dopaminergic pairs (n = 64). Significantly noise-correlated cholinergic-dopaminergic pairs (p < 0.05; 20/64) were always negatively correlated for T1-DANs (10/10) and positively correlated for T2-DANs (10/10).

Importantly, cue-evoked cholinergic and dopaminergic responses largely overlapped in time, whereas feedback responses were temporally separated (Fig. 2B top). Neuronal responses followed a T1-DAN → T2-DAN → BFCN sequence after reward-predicting cues but the reverse, BFCN → T2-DAN → T1-DAN, after rewards and punishments, suggesting that cholinergic-dopaminergic interactions may be organized differently across behavioral epochs (Fig. 2B bottom). Consistent with this temporal organization and the suppression of T1-DAN activity following cholinergic firing, T1-DANs showed suppression after cholinergic cue responses emerged, whereas T2-DANs showed sustained activation that overlapped with cholinergic responses (Fig. 1K and S8B). The opposing response profiles extended to punishment, which inhibited T1-DANs but activated T2-DANs (Fig. 1M and S8B). Finally, T1-DANs lacked the fast reward-response component often observed in T2-DANs (Fig. S8C,D).

In line with these, joint peri-event time histograms revealed distinct dopaminergic cell-type- and task-epoch-dependent modulation of cholinergic-dopaminergic interactions (Fig. S8E-G). Following cues, interactions between BFCNs and T1-DANs consisted of a rapid positive-to-negative correlation sequence, whereas after feedback the delayed inhibitory component predominated. In contrast, BFCN-T2-DAN interactions exhibited strong zero-lag synchrony following cues that became substantially weaker during feedback (Fig. 2C,D and S8H). Cross-correlation analysis of acetylcholine release in the basolateral amygdala and dopamine release in the ventral striatum revealed a positive-to-negative correlation pattern over time, consistent with the opposing interaction patterns observed in the spike-time analysis (Fig. 2E).

If these interaction motifs shape trial-by-trial neuronal variability, they should also be reflected in the co-variation of responses to learned cue-outcome associations. Indeed, fixed cue-evoked noise correlations showed a complete segregation by dopaminergic cell type: all significant BFCN-T1-DAN pairs were negatively correlated, whereas all significant BFCN-T2-DAN pairs were positively correlated (Fig. 2F; r < 0 for 10/10 significant T1-DAN pairs and r > 0 for 10/10 significant T2-DAN pairs). Together, these analyses revealed behaviorally organized synergistic and antagonistic interaction motifs that were segregated by dopaminergic cell type.

### A disynaptic inhibitory circuit underlies cholinergic-dopaminergic antagonism

The delayed inhibition of T1-DANs following cholinergic activity suggested the existence of an inhibitory circuit linking basal forebrain cholinergic and midbrain dopaminergic neurons. As no direct cholinergic HDB→VTA projection has been reported, we hypothesized that this interaction is mediated by an intermediate neuronal population.

A full cross-correlation analysis of HDB and VTA neuronal clusters (Fig. S9) identified a putative GABAergic HDB population that was activated following BFCN firing and, in turn, preceded inhibition of T1-DANs (Fig. 3A), predicting a disynaptic inhibitory pathway (Fig. 3B). Optogenetic stimulation supported this model: activation of BFCNs increased the firing of putative HDB GABAergic neurons while suppressing T1-DAN activity, consistent with a causal disynaptic interaction (Fig. 3C). The optogenetically evoked response latencies further matched the temporal sequence expected for a disynaptic pathway (Fig. S10A,B). BFCN stimulation did not significantly affect T2-DANs (Fig. S10C), nor had dopaminergic stimulation an effect on BFCNs (Fig. S10D).

**Figure 3.**
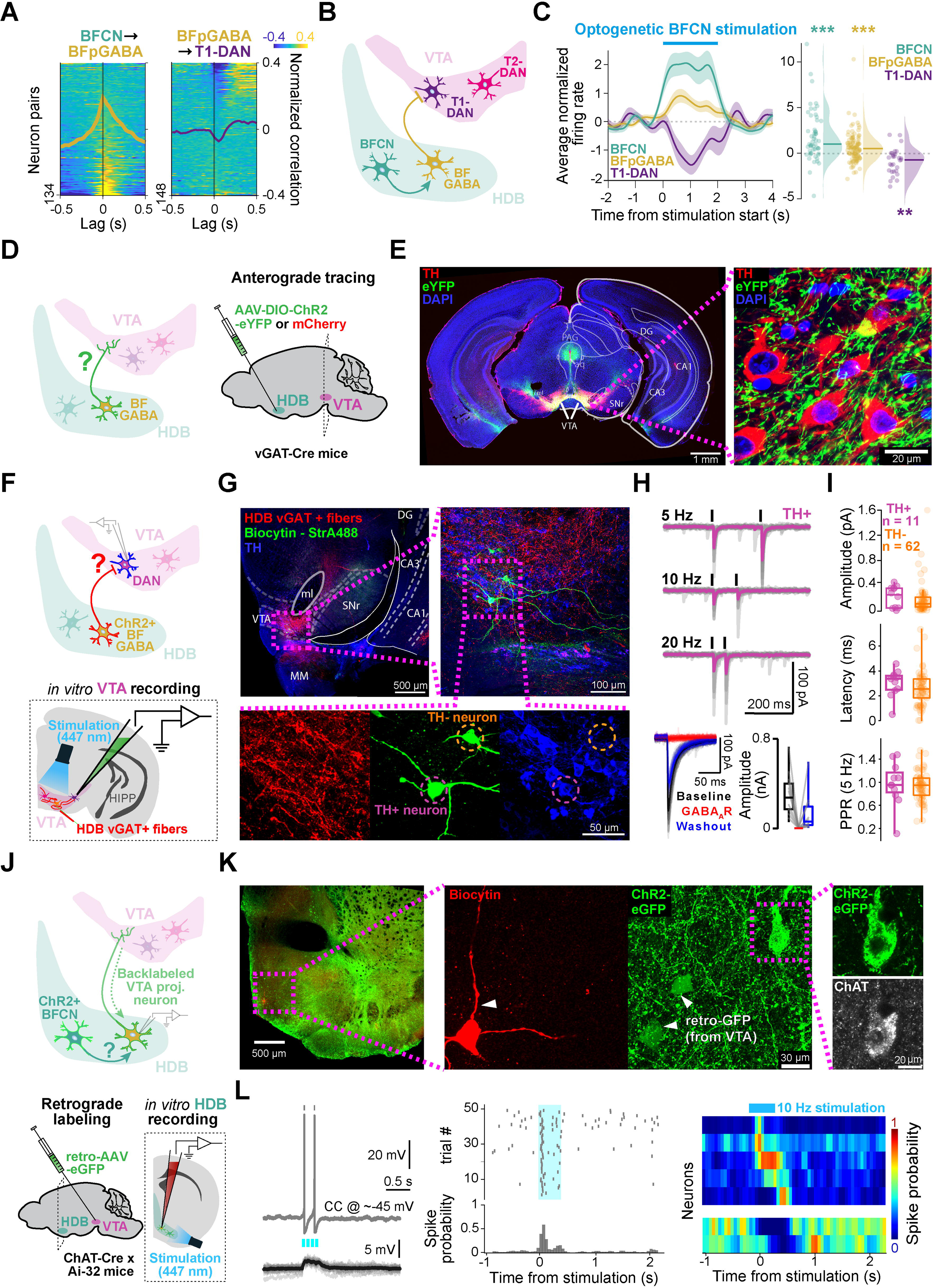
A disynaptic inhibitory pathway from basal forebrain cholinergic to midbrain dopaminergic neurons. **A** Cross-correlograms showing basal forebrain putative GABAergic firing after BFCN spikes (left), and suppression of T1-DAN firing after basal forebrain putative GABAergic spikes (right; rows, pairs of neurons; blue, low cross-correlation; yellow, high cross-correlation; spikes not related to behavioral events (inter-trial intervals) were analyzed; averages are overlaid). **B** Schematic of the proposed circuit. BFCNs activate basal forebrain GABAergic neurons, which inhibit T1-DANs in the VTA. **C** Effects of optogenetic activation of BFCNs on other neurons. Left, average PETHs aligned to photostimulation onset. BFCNs increase their firing rate upon direct stimulation (tagging). Basal forebrain putative GABAergic neurons show increased, while T1-DANs show decreased firing when BFCNs are photostimulated. Error shades, SEM. Right, firing rate changes were statistically significant for BFCNs (p = 4.4 x 10^-6^), basal forebrain putative GABAergic neurons (p = 4.7 x 10^-5^), and T1-DANs (p = 0.0021), but not for T2-DANs (p = 0.1453, two-sided Wilcoxon signed rank test, see Fig. S10C). **D** Schematic of the anterograde tracing experiment of BF GABAergic neurons. The HDB of vGAT-Cre mice were injected with AAV-DIO-ChR2-eYFP or -mCherry. **E** Left: Fluoromicrograph of a coronal section showing TH-expressing dopaminergic neurons (red) and dense fibers of BF GABAergic neurons (eYFP, green) overlapping in the VTA/SNr region. Right: Magnified confocal image from the marked area showing dense GABAergic fibers contacting TH-expressing dopaminergic neurons in the VTA. **F** Schematic of the *in vitro* channelrhodopsin-associated circuit mapping experiment, where VTA neurons were recorded in acute slices while BF GABAergic neurons were optogenetically stimulated. **G** Fluoromicrographs of biocytin-labelled neurons (green; 1/2 TH-expressing) recorded in the VTA. Red, BF GABAergic fibers; blue, TH. **H** Top, IPSCs recorded from an example TH-expressing VTA neuron evoked by photostimulation of BF GABAergic fibers (grey, individual traces; colored, average). Bottom, IPSCs were reversibly blocked by the GABA_A_-antagonist gabazine (10 µM). Left, individual (grey) and average IPSCs under baseline (black), GABA_A_ block (red) and washout (blue) conditions. Right, distribution of IPSC amplitudes in the three conditions (baseline: 248.15 [155.06, 362.12], gabazine: 0.00 [0.00, 0.00], washout: 53.81 [25.92, 176.40] in pA). Box-whisker plots show median, interquartile range and non-outlier range. **I** Distribution of IPSC amplitude, latency and paired-pulse ratio (PPR) at 5 Hz stimulation in TH-expressing (purple) and TH-negative (orange) VTA neurons. IPSC amplitudes in TH-expressing neurons, 288.34 [59.69, 364.38] in pA; in TH-negative neurons, 116.61 [60.34, 219.87] in pA. Latency, TH-expressing neurons, 3.40 [2.53, 3.65] in ms; TH-negative neurons, 2.55 [1.52, 3.35] in ms. Paired-pulse ratios, TH-expressing neurons, 0.96 [0.84, 1.17]; TH-negative neurons, 0.96 [0.79, 1.08]. Box-whisker plots show median, interquartile range and non-outlier range. **J** Schematic of *in vitro* acute slice recordings of VTA-projecting HDB neurons while photostimulating BFCNs locally. ChAT-Cre × Ai32 mice were injected with retro-AAV-eGFP in the VTA. **K** Example of a biocytin-labelled (red) neuron recorded in the HDB, co-expressing eGFP by the retroAAV. Right, note that ChR2-eYFP and retro-eGFP expressions were clearly distinguishable. **L** Light-evoked activity of an example VTA-projecting non-cholinergic HDB neuron upon cholinergic stimulation. Top left, light-evoked spikes in current clamp mode on resting membrane potential. Bottom left, light-evoked postsynaptic excitatory potentials on a hyperpolarized membrane potential below spike threshold. Middle, spike raster (top) and peri-stimulus time histogram (bottom) of light-evoked spikes in the same example neuron. Right, most VTA-projecting non-cholinergic HDB neurons were activated by cholinergic photostimulation. Top, peri-stimulus time histograms aligned to stimulation onset of activated neurons (red, high firing rate; blue, low firing rate). Bottom, the same for inhibited neurons. DG, dentate gyrus, CA1, field CA1 of the hippocampus, CA3, field CA3 of the hippocampus, PAG, periaqueductal gray, aq, aqueduct, MM, medial mammillary nucleus, VTA, ventral tegmental area, SNr, substantia nigra pars reticulata.

To identify the anatomical substrate of the predicted relay, we injected AAV-ChR2-eYFP into the HDB of vGAT-Cre mice (Fig. 3D), revealing a robust GABAergic HDB→VTA projection that innervated both tyrosine hydroxylase (TH)-positive dopaminergic and TH-negative non-dopaminergic VTA neurons (Fig. 3E). Optogenetic activation of HDB vGAT+ axons in acute VTA slices (Fig. 3F,G) evoked inhibitory postsynaptic currents (IPSCs) in both neuronal populations, with approximately half of the recorded TH+ neurons (6/11) exhibiting particularly large-amplitude IPSCs, consistent with the physiological heterogeneity observed in vivo (Fig. 3H,I). Finally, we recorded from retrogradely labeled VTA-projecting HDB neurons in acute slices of ChAT-Cre × Ai32 mice expressing ChR2 in cholinergic neurons (Fig. 3J,K). Photostimulation of local cholinergic neurons activated the majority of VTA-projecting HDB neurons (Fig. 3L), demonstrating that cholinergic neurons can recruit the HDB projection to the VTA. Together, these findings identify a previously unrecognized disynaptic HDB^ACh^ – HDB^GABA^ – VTA^DA^ pathway that provides a circuit mechanism for the delayed cholinergic inhibition of T1-DANs.

### Synergistic cholinergic-dopaminergic interactions support learning

We next investigated how cholinergic-dopaminergic interactions shape learning and trial-by-trial behavioral choices. Novel cue responses of both BFCNs and DANs progressively tracked the evolving behavioral response rate as novel cue-outcome associations were acquired (Fig. 4A and S11A). We therefore asked whether the joint activity of cholinergic and dopaminergic neurons carried complementary information about trial-by-trial behavioral choices. BFCN and DAN activity each predicted behavioral responses, but their predictive information was only partially overlapping: combining cholinergic and dopaminergic activity significantly improved behavioral prediction beyond either population alone (Fig. 4B and S11B). For T2-DANs, this relationship was symmetric: the behavioral associations of BFCNs and T2-DANs were similarly attenuated when accounting for the activity of the other population, consistent with a synergistic relationship. In contrast, for T1-DANs, this relationship was asymmetric: accounting for BFCN activity reduced the T1-DAN–behavior correlation, whereas accounting for T1-DAN activity did not comparably reduce the BFCN–behavior correlation (Fig. 4C and S11C). Consistent with this asymmetry, BFCN activity provided more additional predictive information beyond T1-DANs than T1-DAN activity provided beyond BFCNs (Fig. S11B). At the trial-by-trial timescale, changes in BFCN activity preceded both behavioral adaptation and T1-DAN, but not T2-DAN responses (Fig. 4D and S11D).

**Figure 4.**
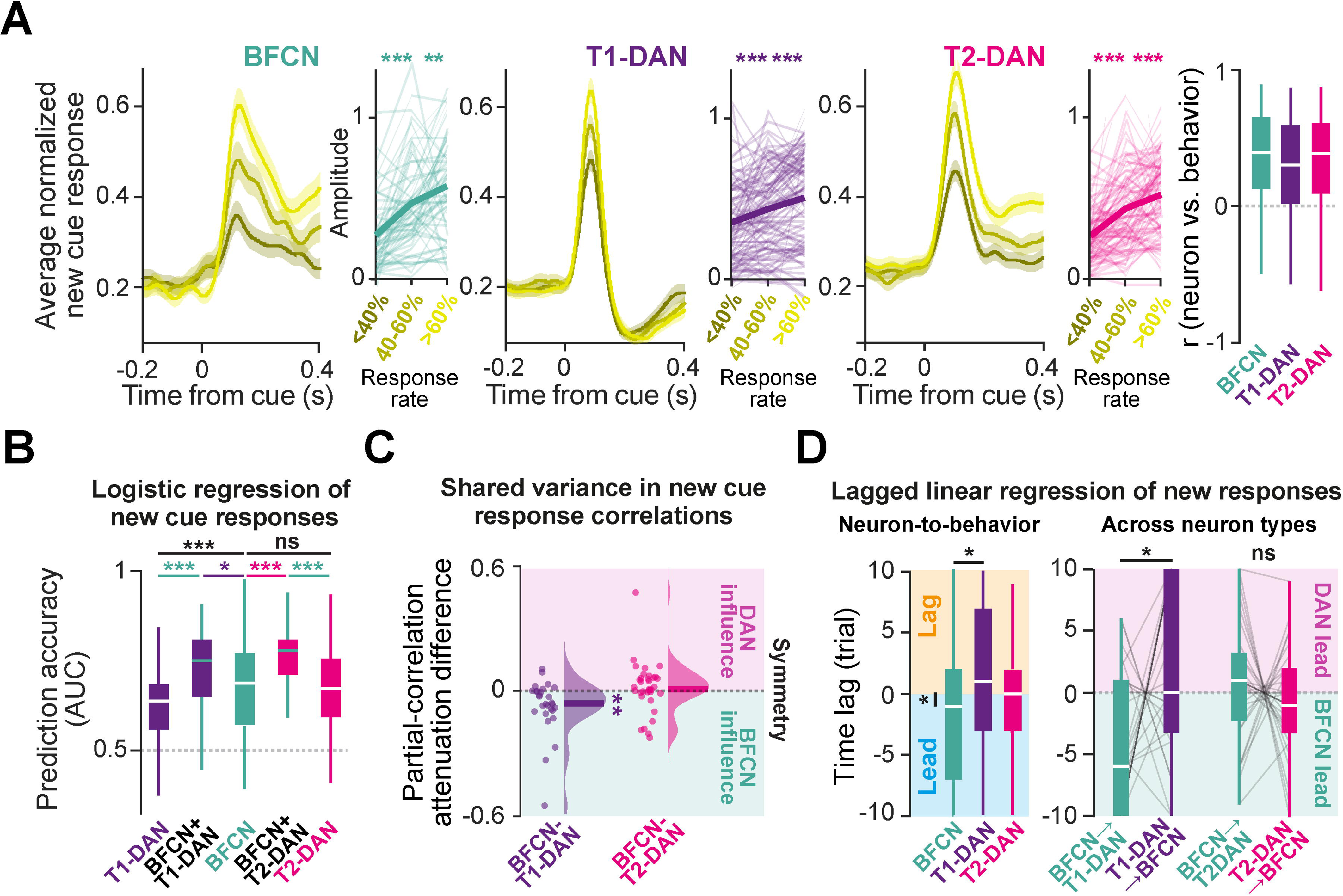
Cholinergic and dopaminergic neurons synergistically predict behavioral choices during learning. **A** Cholinergic and dopaminergic activity robustly represent learning progress indexed by the animal’s responsiveness to the novel cue. Left, average cue aligned normalized PETHs of BFCN, T1-DAN and T2-DAN neurons (error shades, SEM). Insets, statistical comparisons of response amplitudes as a function of mouse responsiveness (BFCN: <40% vs. 40-60%, p = 1.1 × 10^-8^; 40-60% vs. >60%, p = 0.0015; T1-DAN: <40% vs. 40-60%, p = 0.00001; 40-60% vs. >60%, p = 0.000267; T2-DAN: < 40% vs. 40-60%, p = 3.6 × 10^-10^; 40-60% vs. >60%, p = 0.00047; two-sided Wilcoxon signed-rank test). Thin lines, single neurons, thick lines, average. Right, correlations with mouse responsiveness were individually robust and statistically indistinguishable across cell types (two-sided Mann-Whitney U-test). **B** Accuracy in predicting behavioral responses in new-cue trials based on pairs of simultaneously recorded single-neuron responses (logistic regression model; T1-DAN + T1-DAN vs. BFCN + BFCN, p = 1.1 × 10^-5^; T2-DAN + T2-DAN vs. BFCN + BFCN, p = 0.7019). Predictions based on combined neuron types outperformed those based on pairs of neurons from either population alone (BFCN + BFCN vs. BFCN + T1-DAN, p = 0.0499; T1-DAN + T1-DAN vs. BFCN + T1-DAN, p = 1.4 × 10^-5^; BFCN + BFCN vs. BFCN + T2-DAN, p = 6.5× 10^-5^; T2-DAN + T2-DAN vs. BFCN + T2-DAN, p = 2.0 × 10^-7^; two-sided Mann-Whitney U-test). **C** Distribution of differences in partial-correlation attenuation between neuronal response – mouse responsiveness correlations for BFCN-DAN pairs. For T2-DANs, the difference was not significantly different from zero (p = 0.1236), consistent with a symmetric relationship. In contrast, T1-DANs showed an asymmetric relationship: BFCN activity attenuated the DAN–choice correlation more strongly than DAN activity attenuated the BFCN–choice correlation (p = 0.0083; two-sided Wilcoxon signed-rank tests). **D** Left, a linear model was fitted to explain behavioral trends by time-shifted neuronal activity. The majority of BFCNs anticipated mouse responses (p = 0.0445, Wilcoxon signed-rank test, tested against strictly positive values, accounting for the within-trial anticipation of behavioral decisions by neuronal responses) and showed a best fit earlier than T1-DANs (p = 0.0213, two-sided Mann-Whitney U-test). Right, the same model was applied to cross-predict neuronal responses. BFCN responses predicted those of T1-DANs (p = 0.0255), while no such effect was found for T2-DANs (p = 0.191, two-sided Wilcoxon signed-rank test). Box-whisker plots show median, interquartile range and non-outlier range. *, p < 0.05; **, p < 0.01; ***, p < 0.001.

### Cholinergic activity causally shapes dopaminergic prediction-error signaling and learning

These observations led to two testable predictions (Fig. 5A). At reward-predicting cues, dopaminergic responses precede cholinergic activity and thus can escape the disynaptic inhibitory influence identified above (Figs. 2 and 3), so suppressing cholinergic activity should nevertheless reduce dopaminergic cue responses if synergistic cholinergic-dopaminergic interactions reinforce cue-value representations during learning, as suggested by their complementary behavioral information (Fig. 4). Conversely, after punishment, cholinergic activity precedes T1-DAN inhibition, so if cholinergic inhibition contributes to dopaminergic punishment signaling, cholinergic suppression should attenuate this punishment-induced inhibition. These predictions concerned distinct components of dopaminergic signaling—reward anticipation and punishment processing—allowing synergistic and antagonistic cholinergic-dopaminergic interactions to be evaluated independently.

**Figure 5.**
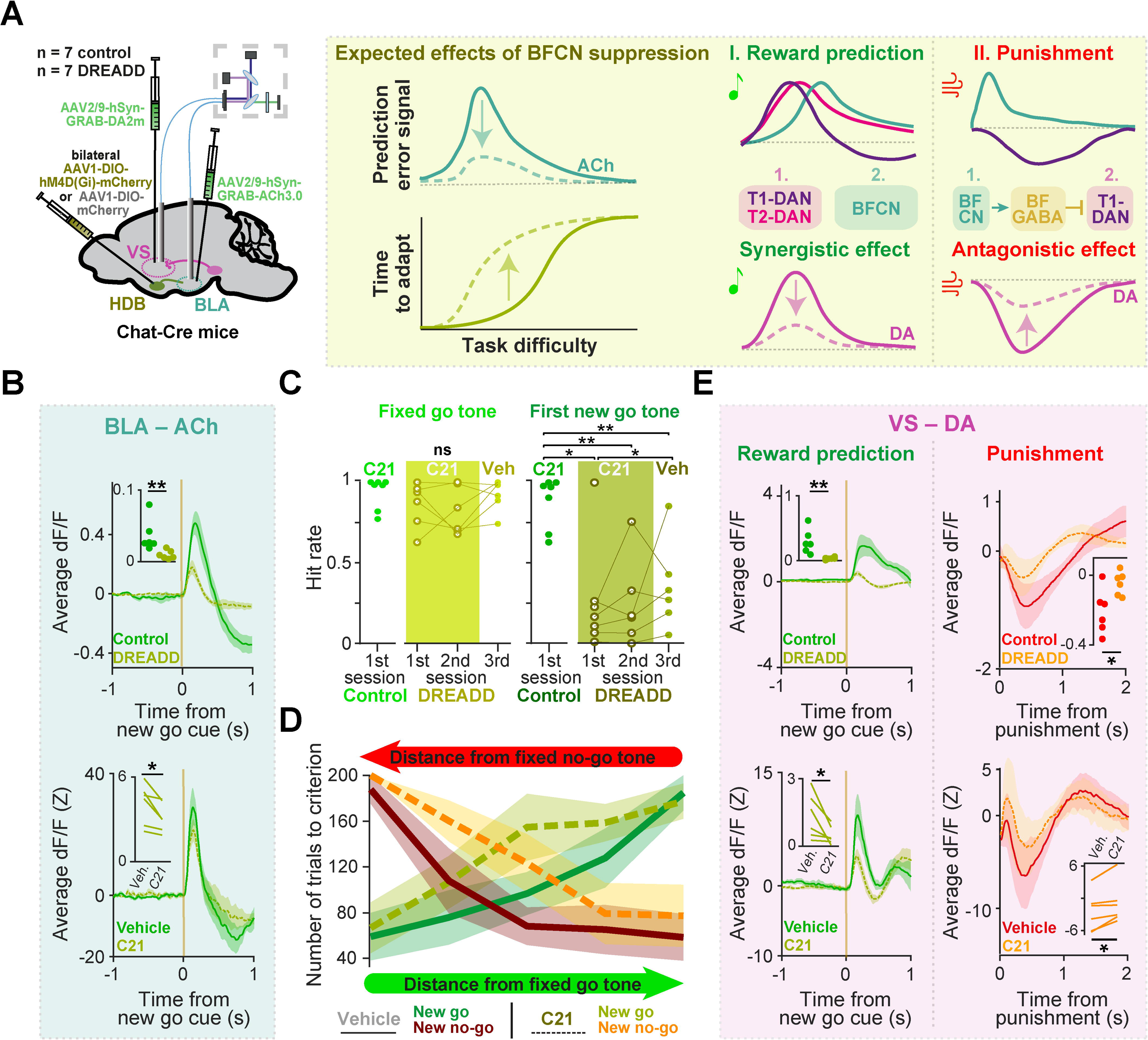
Chemogenetic suppression of basal forebrain cholinergic neurons impairs dopaminergic prediction error coding and learning. **A** Left, schematic of the experiment. ChAT-Cre mice received bilateral injections of AAV1 viral vectors carrying the hM4D(Gi) inhibitory DREADD (n = 7) or control virus (n = 7) in the HDB, and of AAV9 vectors carrying ACh and DA sensors in the BLA and VS, respectively. This allowed us to monitor neuromodulator release using fiber photometry during chemogenetic inhibition of the BFCNs. Right, predicted effects of cholinergic inhibition. First, dopaminergic cue responses precede BFCN responses and may therefore escape the disynaptic inhibitory influence. Cholinergic suppression is predicted to impair reward-prediction coding and behavioral adaptation and, if dopaminergic reward-prediction signals depend synergistically on cholinergic activity (Fig. 4), to reduce DA release following go tones. Second, rapid BFCN activation following punishment precedes T1-DAN suppression, consistent with recruitment of the disynaptic cholinergic pathway (Fig. 3). If this pathway antagonistically shapes dopaminergic punishment signaling, BFCN suppression is predicted to relieve T1-DAN inhibition, resulting in higher DA levels following punishment. **B** Top, average ACh release in the BLA aligned to the novel go cue in control (green) vs. DREADD-expressing (yellow) animals (first session with novel association; error shades, SEM). Inset, cue-evoked ACh release was reduced in DREADD-expressing mice (p = 0.0043; two-sided Mann-Whitney U-test). Bottom, new go tone-evoked ACh release in the BLA of DREADD-expressing mice, Z-scored and averaged across all novel go cues, on days with C21 (yellow) vs. vehicle (green) injection (error shades, SEM). Inset, ACh release was suppressed on days with C21 administration compared to control days with vehicle injection (p = 0.0312, two-sided Wilcoxon signed-rank test). **C** Left, DREADD-expressing mice receiving the C21 DREADD ligand maintained good performance on previously learned fixed go tone-reward associations (compared to control-virus injected mice: 1st new cue session, p = 0.0699; 2nd new cue session, p = 0.1923; 3rd new cue session with vehicle injection, p = 0.366; two-sided Mann-Whitney U-test). Right, at the same time, they failed to acquire the first novel association (compared to control-virus injected mice: 1st new cue session, p = 0.0175; 2nd new cue session, p = 0.0047; two-sided Mann-Whitney U-test). On a control day with vehicle injection (following two C21 sessions), learning deficits still persisted compared to control-virus injected mice (p = 0.0047, two-sided Mann-Whitney U-test), but slightly decrease compared to the first C21 session (p = 0.0312, two-sided Wilcoxon signed-rank test). **D** Psychometric learning curves (see Fig. 1B) comparing sessions of DREADD-expressing mice with C21 vs. vehicle injection revealed pronounced learning deficits at intermediate task difficulty levels. Lines, grand average; error shades, SEM. **E** BFCN suppression had opposing effects on DA release following cues and punishments, consistent with synergistic reward-prediction signaling and antagonistic punishment signaling (see panel A). Top, mean DA release in the VS aligned to the novel go cue (left) and punishment (right) during the first session with a novel association. Control mice are shown in green for go-cue responses and red for punishment responses; DREADD-expressing mice are shown in yellow for go-cue responses and orange for punishment responses. Insets, cue-evoked DA release was reduced (left, p = 0.0022), whereas punishment-evoked DA release was enhanced (right, p = 0.0411) in DREADD-expressing mice (two-sided Mann–Whitney U tests). Bottom, the same responses in DREADD-expressing mice on C21 and vehicle days. Insets, C21 reduced the cue-evoked DA increase (left, p = 0.0312) and attenuated punishment-induced DA suppression (right, p = 0.0312; two-sided Wilcoxon signed-rank tests). Shading, SEM. *p < 0.05; **p < 0.01; ***p < 0.001.

We tested these predictions by chemogenetically suppressing HDB cholinergic activity in ChAT-Cre mice using an inhibitory DREADD (AAV1-DIO-hM4D(Gi)-mCherry injected bilaterally in the HDB; Fig. S12A) while mice performed the psychometric learning task. To determine how cholinergic suppression altered neuromodulatory signaling, we simultaneously monitored ACh release in the BLA and DA release in the VS using GRAB sensors and fiber photometry (Fig. 5A). Chemogenetic suppression of BFCNs markedly reduced ACh release in the BLA compared to both control animals and vehicle sessions (Fig. 5B and S12B), confirming effective suppression of cholinergic signaling.

Chemogenetic suppression of HDB cholinergic neurons profoundly impaired the acquisition of new cue-reward associations. While mice largely retained accurate performance on previously learned fixed tones, administration of the DREADD ligand C21 virtually abolished learning of the first novel association (Fig. 5C), impairing stimulus discrimination and promoting a more conservative response strategy (Fig. S12C–E). Once an initial novel association had been acquired during sessions without C21, mice were able to learn additional associations, although learning remained significantly slower during C21 treatment than on vehicle days (Fig. 5D). These impairments were absent in control animals (Fig. S12F-H). Consistent with a selective deficit in learning rather than sensory discrimination, psychometric functions showed that cholinergic suppression primarily affected novel go and no-go tones of intermediate difficulty (Fig. 5D).

Consistent with the predicted disruption of synergistic cholinergic-dopaminergic interactions, go-cue-evoked DA release was likewise reduced, paralleling the impairment in associative learning (Fig. 5E left). In contrast, the small dopaminergic responses to the fixed no-go cue were preserved or showed a slight increase (Fig. S12I), arguing against a nonspecific reduction in dopamine signaling and instead indicating selective disruption of prediction-error coding.

Cholinergic suppression increased dopamine levels following punishment, thereby attenuating punishment-induced suppression of dopamine release compared to both control animals and vehicle sessions (Fig. 5E right), confirming our second prediction. Consistent with both impaired reward predictions and reduced T1-DAN inhibition (Fig. S12J), dopaminergic responses to less-expected rewards on ‘hard’ trials were also modestly but consistently increased following C21 administration (Fig. S12K). Together, these findings demonstrate that cholinergic neurons are required for the acquisition of novel cue-reward associations and causally shape dopaminergic prediction-error signaling through both synergistic and antagonistic interactions.

## Discussion

We investigated how basal forebrain cholinergic and midbrain dopaminergic projection systems coordinate reinforcement learning. We found that their interactions depended on dopaminergic cell type and task phase, with synergistic and antagonistic relationships emerging across distinct temporal scales and behavioral epochs. During decision making, cholinergic and dopaminergic activity showed synergistic reward-predicting responses that closely tracked, and in the case of cholinergic neurons even anticipated, trial-by-trial behavioral choices. Their activity carried partially complementary information about behavioral choices, such that combining the two populations improved behavioral prediction beyond either population alone. In contrast, BFCNs showed antagonistic interactions with T1-DANs, notably during outcome processing when cholinergic activity preceded T1-DAN inhibition. At the trial-by-trial level, this antagonistic relationship was reflected in negative noise correlations of cue responses. A disynaptic inhibitory pathway from BFCNs to dopaminergic neurons via long-range-projecting GABAergic basal forebrain neurons provided a circuit mechanism for this antagonism. Chemogenetic suppression of BFCNs disrupted dopaminergic reward predictions while attenuating punishment-induced dopamine suppression and impaired acquisition of novel associations, providing causal evidence that cholinergic activity shapes both reward-related and punishment-related dopaminergic signaling.

The cholinergic system was first described using then-novel antibodies against acetylcholinesterase (AChE), the primary degrading enzyme of ACh. However, because DANs also showed strong AChE staining, they were initially included, resulting in a merger of the cholinergic and dopaminergic systems (*36*). This was later resolved using newly developed antibodies against choline acetyltransferase, ACh’s synthetizing enzyme (*37*), and after the two projection systems were precisely delineated (*20*, *38*, *39*), they were rarely considered together.

Instead, researchers sought to interpret cholinergic and dopaminergic functions in orthogonal or complementary terms. Dopamine was associated with learning after the clear demonstration of its strong links to reward (*1*, *40*, *41*), while the cholinergic system was often linked to attention and arousal (*42–48*) or expected uncertainty (*2*). Once DA was established as representing RPE, ACh was proposed to encode learning rate, another key variable of reinforcement learning models (*3*).

However, while these perspectives successfully captured many aspects of ACh dynamics, they tended to overlook strong functional homologies between ACh and DA, some of which have only recently been described. First, lesion and pharmacological studies have shown that both DA and ACh are necessary for associative learning (*49–52*). Second, and relatedly, both systems control postsynaptic plasticity rules, although not necessarily in analogous ways (*53–61*). Third, at the representation level, recent work by us and others has shown that BFCNs, like DANs, respond to rewards and reward-predictive stimuli in a way that can be quantitatively modeled as UPE (*4*, *13*, *62*). This, however, also points to a notable difference: BFCNs show fast and precise positive responses to aversive stimuli, whereas DANs have canonically been described as suppressed, consistent with RPE theory (referred to as T1-DANs in this study). Nevertheless, subsequent studies (e.g. (*25*, *63*)) have reported that a substantial population of DANs is activated by aversive stimuli (referred to as Type 2 DANs), rendering their activity patterns qualitatively similar to UPE-coding cholinergic responses in classical conditioning tasks.

To better understand the unique roles these systems play in broadcasting prediction errors during learning, we developed an operant framework that dissociates rule learning from contingency learning (*64*): mice achieved stable performance on fixed associations while they continually formed flexible new ones in a volatile environment. Moreover, the parametric manipulation of outcome-predicting go cues enabled psychometric analyses of both easy- and hard-to-learn associations, facilitating the detection of subtle changes in learning performance. The coexistence of stable and newly acquired associations allowed us to examine neuronal interactions both moment-to-moment, without the confounding effects of ongoing learning, and across trials as new associations were acquired. We simultaneously recorded BFCNs and DANs while mice were performing this task, overcoming a number of technical challenges including chronic 64-channel dual-bundle tetrode recordings with miniaturized custom-built implants in awake behaving mice, efficient and precise targeting of deep brain nuclei (*65*), and dual optogenetic cell-type identification of sparse populations.

This experimental setting allowed us to test several conditions under which distinct behavioral correlates of prediction errors could emerge (prediction difficulty, learning stage, expectation confidence, temporal uncertainty, and punishment avoidance; Fig. 1, 4, S6, S7 and S11), revealing signal-correlated trial-averaged codes. This argues against orthogonal roles for cholinergic and dopaminergic systems and instead suggests that they act as coordinated components of a distributed reward prediction system that rely on related afferent information about value and reinforcements, supported by shared inputs from the nucleus accumbens, the prefrontal cortex and the lateral hypothalamus (*66–69*). A notable distinction, however, was observed in the sign of punishment responses: T1-DANs exhibited a valence-sensitive, signed prediction error representation, while T2-DANs and BFCNs displayed a salience-sensitive, unsigned prediction error code. This mapping aligns with the long-standing value vs. salience distinction in the dopamine literature (*25*), often associated with separate pathways (such as those targeting ventral vs. posterior striatum (*11*, *70*)) and may extend to cortical cholinergic projections (*20*, *71*).

These observations raise the question of how cholinergic and dopaminergic systems interact, beyond whether they perform overlapping functions. BFCN–T2-DAN interactions were symmetric and centered near zero lag, whereas T1-DANs were asymmetrically suppressed following cholinergic spikes, indicating a directional cholinergic influence on dopaminergic activity. Consistent with this difference, noise correlations also revealed a segregation of the two dopaminergic subtypes: BFCN and T1-DAN responses covaried in opposite directions despite robust positive mean signal correlations, while T2-DAN responses often fluctuated in concert with BFCNs beyond what was explained by their signal correlations (Fig. 2 and S8). Notably, neurons exhibiting signal and noise correlations with opposite signs, sometimes called the ‘sign rule’(*72*), have been shown to carry more stimulus-related information than uncorrelated representations, suggesting that BFCNs may improve stimulus coding in circuits that receive both neuromodulatory inputs (*73*). This was supported by the observation that behavioral choices could be predicted more accurately from combined BFCN and T1-DAN activity than from any single cell type or the combination of the two dopaminergic cell types, with performance statistically indistinguishable from that obtained using combined BFCN and T2-DAN activity, despite T1-DANs alone being less predictive than either BFCNs or T2-DANs alone (Fig. 4B and S11B).

This moment-by-moment BFCN-to-DAN influence was unexpected, because midbrain dopaminergic neurons receive their direct cholinergic input from mesopontine nuclei rather than the basal forebrain (*74*). The latency of DAN responses evoked by optogenetic stimulation of BFCNs suggests that this influence is mediated, at least in part, by a disynaptic pathway, although additional polysynaptic pathways may also contribute. Consistent with this, we identified a disynaptic inhibitory pathway in which basal forebrain cholinergic neurons excite midbrain-projecting basal forebrain inhibitory neurons (*75–77*), which in turn inhibit a population of midbrain dopaminergic neurons (Fig. 3). We propose that this indirect pathway complements previously described direct, subtype-selective mesopontine cholinergic projections – from the pedunculopontine nucleus to T1-DANs and from the laterodorsal tegmentum to T2-DANs (*28*, *69*, *74*) – as well as other non-cholinergic inputs, such as those from the lateral habenula or the ventral pallidum (*28*, *78*). These subtype-selective inputs may be important determinants of the functional heterogeneity of DANs, which are often characterized by their projection targets – T1-like DANs innervate the ventral striatum, while other areas receive mostly T2-like projections (*25*, *79–82*) – or expression profiles (*27*). Notably, while BFCNs were not significantly affected by DAN stimulation, a functional cluster of non-cholinergic basal forebrain neurons – characterized by sustained suppression during reward-predictive cues and reward consumption (Fig. S5, cluster #1) – was inhibited during optogenetic activation of DANs (Fig. S10D). Furthermore, while we found neither dopaminergic projections nor neuronal activation in the HDB upon dopaminergic stimulation, we observed prominent dopaminergic projections in the ventral pallidum, consistent with previous reports (*21*, *83*, *84*), as well as fast and precise activation of some neurons following the optogenetic stimulation of DANs (Fig. S4). Together, these findings suggest that VTA-to-BF feedback pathways also exist, as proposed recently (*85*). Beyond the BF–VTA axis, cholinergic-dopaminergic interactions are well established in the striatum. At striatal terminals, cholinergic interneurons can directly drive DA release via nicotinic receptors (*86–88*), while DA modulates local cholinergic interneurons through D2 receptors (*89*). Recent mesoscale recordings have revealed propagating, largely anti-correlated ACh–DA waves across the striatum, reflecting a spatiotemporally organized sequential neuromodulatory pattern guiding learning and action selection (*90*, *91*).

The temporal organization of these interactions also helps explain the synergistic cholinergic-dopaminergic coding of reward-predicting cues. Cholinergic activity generally preceded dopaminergic activity (particularly T1-DANs) in a number of aspects: (i) when forming new associations, cue-evoked responses of BFCNs anticipated both behavioral responses and T1-DAN activity, which tended to lag behind behavior (Fig. 4D); (ii) cholinergic feedback responses were substantially faster than those of both dopaminergic types (Fig. 2B), likely reflecting a tighter relation to sensory inputs; and (iii) ACh release in the BLA correlated best with subsequent release of DA in the VS (Fig. 2E). Cue-evoked response latencies, however, showed the opposite temporal relationship: dopaminergic responses preceded cholinergic responses (Fig. 2B), allowing DAN activation to escape ACh-mediated disynaptic inhibition. This temporal dissociation may therefore permit reward predictions to be governed primarily by synergistic interactions between the two systems, despite the antagonistic cholinergic influence on T1-DANs observed during outcome processing. Importantly, their synergistic relationship was also evident at the information level: cholinergic and dopaminergic activity each predicted trial-by-trial behavioral choices, but their predictive information was only partially overlapping, such that combining the two populations improved behavioral prediction beyond either population alone. Together, these findings suggest that cholinergic and dopaminergic systems provide complementary information for reward-guided decisions, rather than simply broadcasting redundant signals. More broadly, these temporal relationships suggest a hierarchy in decision-making, progressing from broad, integrative representations of past salient information (T1-DANs), to the current state (T2-DANs), and to prospective, future-oriented planning (ACh, Fig. 4D) – analogous to theta-phase-ordered firing of hippocampal ensembles for past, present, and future locations (*92–95*).

Suppressing cholinergic signaling has been shown to impair sensory learning (*47*, *96*), particularly the encoding of novel associations, while sparing the recall of previously learned associations (*49*, *97*, *98*). However, how this affects the midbrain dopaminergic system, causally related to associative learning (*99*, *100*), has not been investigated. We found that cholinergic inhibition largely abolished the learning of conceptually novel, firstly introduced new associations, made subsequent new associations more difficult to acquire, but only moderately affected performance on previously learned associations (Fig. 5 and S12). Additionally, dopaminergic RPE representations were disrupted, consistent with the predominantly synergistic representation of outcome-predicting stimuli by the two neuromodulatory systems. In parallel, punishment-induced suppression of DA release was reduced, suggesting the causal importance of acetylcholine-mediated disynaptic inhibition of T1-DANs in establishing acetylcholine-dopamine antagonism. Together, these findings demonstrate that cholinergic activity causally shapes dopaminergic prediction-error signaling through both synergistic and antagonistic interactions, linking the two modes of neuromodulatory interaction to learning and punishment processing.

Collectively, these results reframe central cholinergic and dopaminergic signaling as coordinated error-broadcasting systems, consistent with converging evidence for distributed prediction-error codes operating over multiple timescales (*63*, *68*, *101*). The discovery of synergistic trial-by-trial prediction-error codes, together with antagonistic effects mediated by disynaptic acetylcholine-to-dopamine inhibition, bears relevance to disorders involving cholinergic and dopaminergic dysfunction, where mis-calibrated unsigned and signed prediction errors likely interact. Notably, degeneration of BFCNs is a hallmark of cognitive decline in Alzheimer’s disease (*102–107*), while dopaminergic cell loss is associated with impairments of reinforcement-based learning in Parkinson’s disease (*108–114*), likely in a subtype specific manner (*27*, *115–117*). Cross-system effects have also been reported, including cholinergic deficits in Parkinson’s disease dementia (*118–120*), hypercholinergic activity in prodromal Parkinson’s disease without cognitive deficits (*121–123*), selective loss of VTA dopaminergic neurons underlying memory and reward dysfunction in the TG2576 mouse model of Alzheimer’s disease (*124*), and converging human evidence for VTA disconnection and altered dopamine levels in Alzheimer’s disease (*125*, *126*). Our findings suggest that cholinergic and dopaminergic systems should be considered as a joint network rather than independent treatment targets.

## Methods

### Animals

Behavioral experiments were performed in adult male mice (over 2-month-old, 25–30 g) of C57BL/6N genetic background. ChAT-IRES-Cre (n = 2; The Jackson Laboratory, RRID: IMSR_JAX:006410), DAT-IRES-Cre (n = 1; The Jackson Laboratory, RRID:IMSR_JAX:006660) or ChAT-Cre x DAT-Cre (n = 12; obtained by crossing the former two lines) mice were used for optogenetic tagging experiments. Wild type C57BL/6N mice were used for fiber photometry experiments (n = 10). Optogenetic circuit mapping was performed in vGAT-IRES-Cre mice (n = 15; The Jackson Laboratory, RRID: IMSR_JAX:016962) to test HDB to VTA connection. To characterize cholinergic responses onto VTA-projecting HDB neurons, we crossed ChAT-IRES-Cre with Ai32 (n = 4; Jackson laboratories, RRID: IMSR_JAX:012569) mice. Anterograde tracing of VTA-to-HDB projections was performed in DAT-IRES-Cre (n = 6), vGluT2-IRES-Cre (n = 2; The Jackson Laboratory, RRID:IMSR_JAX: 016963) and vGAT-IRES-Cre (n = 4) mice. Chemogenetic suppression of BFCNs combined with behavioral training and fiber photometry recordings was performed in ChAT-IRES-Cre mice (n = 14, 7/14 mice served as controls). Animals were housed individually in 36 × 20 × 15-cm cages under a standard 12-h light–dark cycle (lights on at 8 a.m.) with food available ad libitum. Mice were water-restricted during behavioral training. Temperature and humidity were kept at 21 ± 1 °C and 50–60%, respectively.

All experiments were approved by the Institutional Animal Care and Use Committee and the Committee for Scientific Ethics of Animal Research of the National Food Chain Safety Office (PE/EA/675-4/2016, PE/EA/1212-5/2017, PE/EA/864-7/2019 and PE/EA/1003-7/2021), and were performed according to the guidelines of the institutional ethical code and the Hungarian Act of Animal Care and Experimentation (1998, XXVIII, section 243/1998, renewed in 40/2013) in accordance with the European Directive 86/609/CEE and 2010/63/EU.

### Surgical procedures

Animals were anesthetized with an intraperitoneal injection of a ketamine-xylazine mixture (83 mg/kg ketamine and 17 mg/kg xylazine in 0.9% NaCl). The scalp was shaved and disinfected with Betadine, and Lidocaine was applied subcutaneously to provide local anesthesia. The eyes were protected using an ophthalmic lubricant (Corneregel eye gel, Bausch & Lomb). Mice were placed in a stereotaxic frame (David Kopf Instruments), the skin was incised, the skull was cleaned and leveled using Bregma, Lambda and a pair of lateral points equidistant from the sagittal suture. Craniotomies were opened above the target areas and glass pipettes pulled from borosilicate capillaries, broken to 20–30-μm tip diameter, were lowered stereotaxically. Adeno-associated virus vectors were injected using a MicroSyringe Pump Controller (World Precision Instruments). In the in vivo electrophysiology experiments we injected the viral vector AAV2/5-EF1a-Dio-hChR2(H134R)-eYFP-WPRE-hGH (300 nl; titer, 7.7 × 10^12^ vg/ml; Penn Vector Core) in the HDB (+0.74 mm anterior and 0.6 mm lateral from Bregma, −5.0 and −4.7 mm ventral to the surface) and the VTA (−3.1 mm posterior and 0.6 mm lateral to Bregma, −4.4 and −4.0 mm ventral to the surface). In the fiber photometry experiments we injected the acetylcholine sensor AAV9-hSyn-GRAB-ACh3.0 (300 nl; titer, ≥ 1×10¹³ vg/ml, BrainVTA) to the BLA (−1.2 mm posterior and 2.8 mm lateral to Bregma, −3.7 mm ventral to the surface) and the dopamine sensor AAV9-hSyn-GRAB-DA2m (300 nl; titer, ≥ 2×10^12^ vg/ml; BrainVTA) to the VS (1.0 mm anterior and 1.1 mm lateral to Bregma, −4.0 mm ventral to surface). In the chemogenetic experiments, in addition to the GRAB sensors, the inhibitory DREADD AAV1-hSyn-DIO-hM4D(Gi)-mCherry (240 nl in each hemisphere; titer, ≥ 7×10¹² vg/ml, Addgene) or the control virus ssAAV1/2-hSyn-DIO-mCherry (240 nl in each hemisphere; titer, 5.4 x 10^12^ vg/ml; Viral Vector Facility, Universität Zürich) were injected to the HDB bilaterally. For the anterograde tracing of VTA-to-HDB projections, mice received bilateral injections of AAV2/8-CAG-Flex-eGFP (30 nl in each hemisphere; titer, ≥ 1 × 10¹³ vg/mL; Addgene) into the VTA. In anterograde optogenetic circuit mapping from the HDB to the VTA, animals received injections of either ssAAV9/2-hEF1α-DIO-hChR2(H134R)-mCherry-WPRE-hGHp (60 nl; physical titer, 6.0 × 10¹² vg/mL; VVF, Zürich) or AAV2/5-EF1a-DIO.hChR2(H134R)-eYFP-WPRE-hGH (60 nl; titer, ≥ 1 × 10¹³ vg/mL; Addgene) into the HDB. For optogenetic circuit mapping of cholinergic modulation of VTA-projecting HDB neurons, ssAAV-retro/2-CAG-EGFP-WPRE-SV40p(A) (300 nl; titer, 5.3 × 10¹² vg/mL; VVF, Zürich) was injected into the VTA.

For chronic in vivo electrophysiological recordings, we designed a custom-built microdrive that carried two independently movable implants, each consisting of eight tetrode electrodes (nichrome, 12.7 µm diameter; Sandvik) connected to an Omnetics connector, and a 50-µm-core optic fiber (outer diameter, 65 ± 2 µm; Laser Components GmbH, Olching, Germany), enabling parallel implantation into the HDB and VTA. Microdrive screws (M0.6 stainless steel flat head screw, 12 mm length, 160 µm thread pitch; Easterntec) provided controlled advancement of the tetrodes and the optic fiber with ∼20 µm precision. For gold-plating, polyethylene glycol (PEG) was dissolved in deionized water to a concentration of 1 g/l, and 1.125 ml of this PEG solution was mixed with 0.375 ml of gold-plating solution (Neuralynx), following (*127*). Electrodes were gold-plated to impedances of 30–100 kΩ measured at 1 kHz using NanoZ (Cambridge NeuroTech). Before implantation, a fluorescent dye (DiI, Invitrogen LSV22885) was applied to the tips of the tetrode bundles to enable later histological track reconstruction. Two additional holes were drilled in the occipital plate for the ground and reference wires. The implants were lowered stereotaxically to the top of the target areas, the craniotomies were covered by low viscosity silicone elastomer sealant to protect the brain surface, and the implant was secured using dental adhesive (C&B Metabond, Parkell) and acrylic resin (Jet Denture, Lang Dental). For fiber photometry experiments, optic fibers (400 μm core, ceramic ferrules) were bilaterally implanted targeting the BLA and the VS, 0.1 mm above the injection sites. The fibers were fixed to the skull using Super-Bond (Sun Medical Co.) and dental cement. After surgery, Buprenorphine (Bupaq, 0.3 mg ml⁻¹, Richter Pharma AG) was used to provide postoperative analgesia and gentamycin as a local antibiotic. The surgery was followed by a 10-day recovery period before behavioral training began.

### Behavioral training

Mice were trained on a psychometric operant learning task according to a head-fixed auditory go no-go discrimination paradigm. We used a custom-built setup equipped with a head-fixing apparatus, a white LED used to indicate trial-start, two speakers delivering calibrated auditory sensory cues, a lickometer to detect behavioral responses, a water delivery port, and an air-puff delivery system (*128*). We used the open source Bpod behavioral control system (Sanworks LLC, US) for operating the task. Mice were water-scheduled to achieve 85-90% of normal body weight. Drops of water (3 µl) were used as water reward after correct go responses and puffs of air directed to the animals’ face were used as punishment (200 ms duration, 15 psi, reported as aversive for mice (*12*, *129*)).

The start of each trial was signaled by turning off a white LED. This was followed by a variable foreperiod drawn from an exponential distribution (mean 1.3 s, truncated between 0.3–4 s) to avoid temporal expectation of the cue stimuli (*130*, *131*). If the animal licked during the foreperiod, the trial was reset and the LED was turned on again. After the foreperiod, a cue tone was played. Initially, animals were trained exclusively on a fixed reward-predicting tone (pure tone of 4 kHz frequency, 0.6 s duration, 50 dB sound pressure level). If mice broke the photobeam of the lickometer with their tongues (lick response) in a 30 s response window after go tone onset, they received water reward from a waterspout placed close to their snout. After establishing a reliable tone–reward association (≥50 hits with response times within 0.6 s from tone onset), the response window was automatically shortened to its final duration (0.6 s). Once animals reached stable performance (hit rate ≥70%), a well distinguishable (6 kHz higher) fixed punishment-predicting no-go tone was introduced in 50% of the trials. If mice licked after the no-go tone, they received a facial air puff as punishment. The cue stimulus was stopped when mice responded. Not licking after a go tone was considered a missed trial, while withholding lick responses after a no-go tone was considered a correct rejection. The feedback was delivered in hit and false alarm trials after a normally distributed delay with respect to mouse response (mean, 0.2 s; standard deviation, 0.03 s; truncated between 0.1–0.3 s) to temporally dissociate motor responses from reinforcement. After outcome delivery, the LED was turned on again, and a new trial began if mice stopped licking for at least 1.5 s. In the photometry experiments, cue duration and feedback delay were increased to 1 s and 0.6 s, respectively, to accommodate slower sensor signals compared to spiking data. However, the original parameters were kept in the chemogenetic suppression experiments for better behavioral comparability.

The fixed associations were kept constant throughout the entire training. When mice reached 70% performance in both fixed associations, a new go association was introduced at mid frequency between the fixed cues. When mice reached ≥70% correct responses in all three trial types (averaged over the last 50 trials from each type, or, if less available, over all trials after a minimum of 33 trials), the novel association was replaced randomly with either a new go or no-go tone, with its frequency randomly drawn from the intermediate integer frequencies between the two fixed tones, avoiding immediate repetition. Novel cue-outcome pairings always replaced half of the fixed associations of the same valence, in order to keep the ratio of go and no-go trials 1:1. If mice did not reach the 70% criterion within 200 new tone trials, learning of the new association was considered unsuccessful, and a new cue-outcome pair was introduced at the start of the next session.

### Fiber photometry

Simultaneous fluorescent sensor signals (ACh in the BLA and DA in the VS) were measured with a dual fiber photometry setup (Doric Neuroscience). Two LED light sources (465 nm and 405 nm) matching the maximum excitation and isosbestic wave lengths of the sensors were amplitude-modulated by the command voltages of a two-channel LED driver (LEDD_2, Doric Neuroscience) — with the 465-nm and 405-nm LEDs driven at 208 Hz and 572 Hz, respectively—and directed into fluorescent Mini Cubes (iFMC4, Doric Neuroscience). In two animals, only a single Mini Cube was used, and ACh and DA signals were recorded in alternating sessions. Light was delivered through 400-µm patch-cord fibers and coupled to the implanted optical fibers during training sessions. The same fibers were used to collect the bilaterally emitted fluorescence, which was detected by the 500–550 nm photodetectors integrated in the Mini Cubes. Fluorescence signals were sampled at ∼12 kHz and demodulated offline.

### In vivo implant localization

Implant position was verified in vivo (*65*). Preoperative micro-CT and T1-weighted MRI images were acquired under isoflurane anesthesia using a NanoX-CT cone-beam CT system (Mediso Medical Imaging Systems) equipped with an 8-W power X-ray tube (tube voltage, 45 kVp; magnification, 1.36; exposure time, 500 ms; number of projections, 180) and a nanoScan PET/MRI (Mediso Medical Imaging Systems) with a permanent magnetic field of 1 T and with a 450 mT m^−1^ gradient system (3D gradient echo sequence using 8 excitations, T_R_ = 15-ms repetition, T_E_ = 2.2-ms echo times and 25° flip angle with resolution set to 0.28 mm). 4-12 days after the surgery, post-operative micro-CT imaging was performed (tube voltage, 45 kVp; magnification, 2.47; exposure time, 500 ms; number of projections, 180). Nucline 2.01 (Mediso Medical Imaging Systems) was used for CT image reconstruction with a filtered back projection using a Butterworth filter. The voxel size was isotropically set to 35 µm or 19 µm for 1.36 and 2.47 magnifications, respectively. Once all images were acquired, they were co-registered in the VivoQuant (version: 1.22; inviCRO) software based on anatomical landmarks. Finally, a mouse brain atlas (*132*) was fitted based on the pre-operative MRI images and the Bregma and Lambda points identified on the pre-operative CT images to localize the position of the implant tip, visible on the post-operative CT images.

### Histological track reconstruction and anterograde tracing

After completing the experiments or 4 weeks after viral injection for tracing, mice were deeply anesthetized with a ketamine–xylazine mixture. In animals implanted with tetrode drives, the final electrode position was marked for subsequent histological reconstruction by producing a small electrolytic lesion (40 µA for 5 s through one or two tetrode channels; IonFlow Bipolar, Supertech Instruments, Pécs, Hungary). Mice were then transcardially perfused with 0.1 M phosphate-buffered saline (PBS) for 1 min, followed by 4% paraformaldehyde (PFA) in PBS for 20 min. Implants were carefully removed, and brains were extracted and post-fixed in PFA for 24 h at 4 °C.

Coronal sections (50 µm) were cut using a Vibratome (VT1200S, Leica, Wetzlar, Germany). For the animals that underwent DREADD injections, sections were stained with 4′,6-diamidino-2-phenylindole (DAPI; 10 mg ml⁻¹; Merck Millipore, Burlington, MA, USA). Slices were mounted on microscope slides, coverslipped with Vectashield mounting medium (Vector Labs, Burlingame, CA, USA).

For track reconstruction, a Nikon Eclipse Ni fluorescence microscope equipped with a Nikon DS-Fi3 camera was used to acquire bright-field, dark-field, and fluorescence images. Images were processed to reconstruct implant trajectories. Corresponding planes from a mouse brain atlas (*133*) were aligned to each section using linear scaling and rotation, guided by anatomical landmarks visible in bright-field and dark-field images. Atlas overlays were transferred onto fluorescence images that revealed virally transfected ChR2–eYFP and Dil-labeled tetrode tracks or sensor expression. The coordinates of tetrode tips and the orientations of their trajectories were determined from the Dil-labeled paths and the locations of electrolytic lesions identified in bright-field and dark-field images. For the movable tetrode implants, the moving distance was logged on each recording day, and the corresponding coordinate of the implant was calculated based on the reconstructed tip position and trajectory. For anterograde tracing, fluorescent images were acquired using a Nikon A1R confocal laser scanning microscope at 10× magnification and processed similarly. Axonal projection density was quantified by calculating the mean pixel intensity within each target region.

### In vivo electrophysiology

In vivo extracellular electrophysiology data were acquired using the Open Ephys data acquisition system. Two 32-channel Intan headstages (RHD2132) were connected to the Omnetics connectors on the custom-built microdrive, and signals were transmitted via digital SPI cables (Intan) to the Open Ephys acquisition board, which received TTLs from the BPod behavior control unit and the Pulse Pal stimulus generator for synchronization. Data were digitized at 30 kHz and recorded using the Open Ephys software. Implants were advanced in 20-80 μm steps based on single unit yield and the occurrence of light-evoked potentials.

### Optogenetics

The microdrive for in vivo electrophysiology recordings held optic fibers with FC connectors (Precision Fiber Products). These were connected to FC-APC patch cords during recordings. Before and after each training session, BFCNs and DANs were optogenetically identified separately by delivering 1 ms laser pulses (473 nm, Sanctity) at 20 Hz for 2 s followed by a 3 s inter-pulse interval driven by a stimulus generator (Pulse Pal, Sanworks), repeated 24 times. Light-evoked spikes and potential artifacts were monitored online using the OPETH plugin (SCR_018022, (*18*)) for Open Ephys. Whenever photoelectric artifacts or population spikes were detected, laser intensity was adjusted to prevent masking of individual action potentials.

### Acute in vitro slice preparation

Mice were decapitated under deep isoflurane anesthesia, and the brains were rapidly removed and placed in ice-cold cutting solution, pre-carbogenated (95% O₂–5% CO₂) for at least 30 minutes before use. The cutting solution consisted of (in mM): 205 sucrose, 2.5 KCl, 26 NaHCO₃, 0.5 CaCl₂, 5 MgCl₂, 1.25 NaH₂PO₄, and 10 glucose. Coronal slices, 300 μm thick, were prepared using a Vibratome (Leica VT1200S). Following acute slice preparation, slices were transferred to an interface-type holding chamber for at least one hour of recovery. This chamber contained ACSF solution maintained at 35 °C, which gradually cooled to room temperature. The ACSF solution consisted of (in mM): 126 NaCl, 2.5 KCl, 26 NaHCO₃, 2 CaCl₂, 2 MgCl₂, 1.25 NaH₂PO₄, and 10 glucose, saturated with carbogen gas as described above.

### In vitro electrophysiology recordings

Recordings were performed under visual guidance using Nikon Eclipse FN1 microscope with infrared differential interference contrast (DIC) optics. The flow rate of the ACSF was 4–5 ml/min at 30–32°C (Supertech Instruments, Pécs, Hungary). Patch pipettes were pulled from borosilicate capillaries (with inner filament, thin-walled, outer diameter (OD) 1.5) with a PC-10 puller (Narishige, Tokyo, Japan). Pipette resistances were 3–6 MΩ when filled with intrapipette solution. The composition of the intracellular pipette solution used for VTA recordings was as follows (in mM): 54 d-gluconic acid potassium salt, 4 NaCl, 56 KCl, 20 Hepes, 0.1 EGTA, 10 phosphocreatine di(tris) salt, 2 ATP magnesium salt and 0.3 GTP sodium salt; with 0.2 % biocytin; adjusted to pH 7.3 using KOH and with osmolarity of ∼295 mOsm/l. The composition of the intracellular pipette solution used for HDB recordings was as follows (in mM): 110 d-gluconic acid potassium salt, 4 KCl, 20 Hepes, 0.1 EGTA, 10 phosphocreatine di(tris) salt, 2 ATP magnesium salt and 0.3 GTP sodium salt; with 0.2 % biocytin; adjusted to pH 7.3 using KOH with osmolarity of 290 mOsm/l. Recordings were performed with a Multiclamp 700B amplifier (Molecular Devices, San Jose, US), digitized at 10 or 20 kHz with Digidata analog-digital interface (Molecular Devices), and recorded with pClamp11 Software suite (Molecular Devices). Retrogradely back-labeled GABAergic neurons expressing GFP in the HDB, or their eYFP expressing axons in the VTA were visualized with the aid of LED light sources (Prizmatix Ltd., Holon, Israel) integrated into the optical light path of the microscope and detected with a CCD camera (Andor Zyla). In case of characterizing HDB to VTA GABAergic connection, the bath solution contained 20 μM NBQX and 50 μM AP5 to block glutamatergic excitatory currents.

### Immunohistochemical identification of in vitro recorded GABAergic HDB and TH+ VTA cells

After acute slice electrophysiology experiments, brain sections were fixed overnight in 4% PFA. Sections were extensively washed in 0.1M PB and TBS and blocked in 1% human serum albumin (HSA; Sigma-Aldrich) solution for 1 h. Then, sections were incubated in primary antibody against TH (Cat#213106, SYSY, 1:500, raised in chicken), GFP (Cat#ab5450, Abcam, 1:2000, raised in goat) or mCherry (Biovision, Cat#5993-100, 1:1000, raised in rabbit) for 48-60 hours. This step was followed by thorough rinse with TBS (3 × 10 minutes) and overnight incubation with a mixture of anti-chicken DyLight-405 (Jackson Immunoresearch, Cat#103-475-155, 1:500, raised in goat) anti-rabbit Alexa-594 (Thermo Fisher Scientific, Cat#A21207, 1:500) or anti-goat Alexa-488 secondary antibody (Cat#705-545-147 Jackson Immunoresearch, 1:500, raised in donkey) and streptavidin-A488 (Invitrogen, Cat#S11223, 1:1000) or streptavidin-A594 (Invitrogen, Cat#11227, 1:1000). We used 0.1% Triton-X detergent through every incubation step due to the thickness of the brain section. Finally, sections were washed in TBS and PB, mounted on microscopy slides, covered with Vectashield (Vector Laboratories Inc, US) and imaged with a Nikon A1R confocal laser scanning microscope.

### Chemogenetics

For the chemogenetic experiments, we used C21 (Merck) as the ligand at a dose of 1 mg/kg (stock concentration: 0.2 mg/ml dissolved in physiological saline) following the dosing guidelines described in (*134*). The compound was administered subcutaneously 30 minutes before the training sessions, separated by at least 24 hours. Mice were habituated to injections during the pre-training handling period. Initially, animals underwent needle insertions without injection. From the fourth day of handling, they were injected with saline until 70% performance on both fixed associations was reached. When the first new association was introduced, animal received C21 treatment for two consecutive sessions, followed by saline sessions until the first new association was not formed. After this, C21 administration followed an alternating schedule: two consecutive days of C21 injection were followed by two days of saline injections serving as self-control.

### Data analysis

Data analysis was carried out using custom written Matlab (R2016a, MathWorks) or Python 3 codes.

### Fluorescent sensor signals

Fiber photometry data were preprocessed following the procedure described by (*135*). First, a 20 Hz low-pass Butterworth filter was used to remove high-frequency noise. Second, the isosbestic 405 nm signal (F_405_) was fitted to the neuromodulator-dependent 465 nm (F_465_) signal using a least-squares linear regression to match baseline intensities. Third, this fitted isosbestic signal (F_405, fitted_) was used to normalize F_465_:

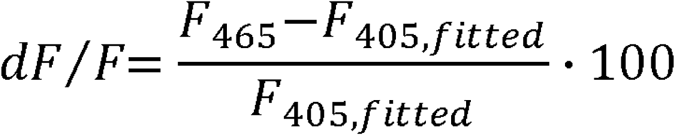

This normalization corrected for motion artifacts and autofluorescence. Slow baseline drifts were removed using a 0.2 Hz high-pass Butterworth filter. The resulting ΔF/F signal was aligned to cue and feedback events, and averaged across trials, sessions and animals. Event-evoked responses were quantified as the area under the curve (AUC) between the half-maximum/half-minimum locations around the peak/trough responses, with the dF/F value at the first half-maximum/half-minimum point serving as the baseline. When comparing the effect of C21 to control days across animals in the chemogenetic experiments, the dF/F signals were Z-scored session-by-session relative to the pre-cue baseline period (2 s prior to cue onset).

### Spike sorting

Tetrode signals were digitally referenced to a common average reference and band-pass filtered between 700 and 7000 Hz using a zero-phase Butterworth filter. Spike detection was performed with a 750 µs censoring period. Action potentials were manually sorted into putative single units offline using MClust 3.5 (A. D. Redish), clustering action potentials in a feature space spanned by the waveform energy and the first principal component features of the waveforms. Spike waveforms were visually inspected to confirm biologically plausible shapes and amplitudes and to exclude noise artifacts or multiunit activity. Furthermore, autocorrelograms were examined for violations of the refractory period, and cross-correlograms between simultaneously recorded units were inspected to exclude duplicate or overlapping units. Only well-isolated putative single units with isolation distance > 20 and L-ratio < 0.15 were included (*136*).

### Optogenetic tagging

Single unit firing following laser pulses during optogenetic stimulation was statistically assessed with the Stimulus-Associated Spike Latency Test (SALT) (*16*). Spike shape correlations (R) between laser-evoked (defined as the spikes occurring within the time window corresponding to the full width at half height of the laser-evoked PETH peak) and spontaneous spikes were calculated (*24*). Neurons with p < 0.01 and R > 0.85 were considered optogenetically identified neurons (n = 90), i.e. BFCNs in the basal forebrain and DANs in the VTA. The median correlation coefficient of all identified neurons was R = 0.99. Consistent with direct light activation, all neurons showed light-evoked spikes within a few milliseconds of light onset with small jitter (Fig. S4D).

### Clustering of single-unit response profiles

Single units were clustered based on Z-score-normalized hit trial PETHs aligned to fixed go cues and rewards and false alarm trial PETHs aligned to punishments. Time windows of the PETHs were chosen to best capture the time scale of cholinergic and dopaminergic responses (from cue onset until feedback time, truncated at 0.4 s; from feedback, 0 to 0.05 s for HDB and 0 to 0.2 s for VTA neurons). Principal component analysis (PCA) was performed separately on hit and false alarm PETHs for dimensionality reduction. The number of components retained was determined using the elbow method (see (*22*)), which indicated that the first two principal components from hit trials and the first principal component from false alarm trials accounted for most of the variance in the dataset (Fig. S5B). Neurons were subsequently clustered using k-means in the three-dimensional space spanned by these principal components yielding five distinct clusters for each brain region (Fig. S5C). The number of clusters was determined using the elbow method (Fig. S5B). In sessions with fewer than five punishments, the punishment-aligned PETHs could not be reliably computed. In these sessions, 13 VTA neurons (7 optotagged) and 8 HDB neurons (3 optotagged) showing clear BFCN- or DAN-like cue- and reward-aligned activity were manually classified. In these cases, Type 1 and Type 2 DANs were distinguished based on the presence of an undershoot versus sustained activity following the initial cue response, as well as punishment-aligned raster plots from a few trials when available.

### Trial-averaged comparison of single unit activity

We calculated event-aligned raster plots and peri-event time histograms of spike times and lick response times (PETHs) aligned to cue onset, feedback delivery and photostimulation start. PETHs were smoothed using Gaussian kernels selected according to the temporal dynamics of the responses. Behavioral event-aligned PETHs were generally smoothed with a 20 ms kernel. To better resolve sharp feedback responses, feedback response signal correlations (Fig. 1L-M, S6B-D, S7 and S11A) were examined with a narrower 10 ms kernel. Photostimulation-onset-aligned PETHs were smoothed using either a broad 200 ms kernel to illustrate sustained effects (Fig. 3C and Fig. S10C–D) or a narrow 1 ms kernel to resolve pulse-locked responses (Fig. 1I, S4G and Fig. S10A). To calculate average PETHs, individual PETHs were Z-score-normalized using the mean and standard deviation of the pre-cue or pre-stimulus activity. In the analysis of the sustained effect of optogenetic stimulation (Fig. 3C and S10), only animals with tagged neurons were included to ensure that the optogenetic stimulation was efficient.

To examine signal correlations (Fig. 1N, 4A, S6 and S11A), trials were grouped by new cue difficulty (frequency distance from fixed cues), new cue responsiveness (<40%, 40–60%, >60%), fixed go cue reaction time (slowest vs. fastest tercile), and easy reward or punishment delays. Reward delays were binned as short (lowest sixth), long (highest sixth), or average (remaining trials), approximately reflecting values below, above, or within ±1 SD of the Gaussian-distributed delays. Punishment delays, with much fewer trials, were similarly binned as short (lowest fifth), long (highest fifth), or average (remaining trials). Cue-evoked PETHs were normalized to the peak fixed go cue response, whereas feedback-evoked PETHs were normalized to the mean peak response to the corresponding feedback across all trials, irrespective of grouping. Positive response amplitudes were quantified as the maximum value within a 200 ms response window relative to pre-event baseline; suppression of activity of T1-DANs after punishments was quantified as the minimum value. To exclude potential effects of recording instability, neurons with robustly different (Mann–Whitney U-test, p < 0.001) pre-cue firing rate in any of the novel cues compared to fixed go cue trials were omitted from the signal correlation analysis.

To assess unimodal and bimodal dopaminergic reward responses (Fig. S8C,D), reward-aligned PETHs were fitted with a single Gaussian function (unimodal) or with the sum of two Gaussian functions (bimodal). The widths of the Gaussians were constrained to be at least as large as the PETH smoothing kernel. For bimodal fits, the peaks were constrained to be separated by at least four kernel widths, and the smaller peak amplitude had to reach at least one-third of the larger peak. Fits were obtained by least-squares optimization, and model preference was assessed using the Bayesian Information Criterion (BIC).

To quantify the temporal coupling between simultaneously recorded cholinergic and dopaminergic responses, peri-event time histograms of each neuron were z-scored to the pre-event baseline and cross-covariance functions were computed within a 0.4 s response window following cue, reward and punishment (Fig. S8B). The component attributable to the product of the two mean responses was subtracted, and the result was normalized by the product of the response standard deviations, yielding a signal-independent measure of co-fluctuation. Temporal overlap between all cholinergic and dopaminergic neuron responses was quantified by the Jaccard index (i.e. intersection over union) of their response windows, defined as the full width at half-prominence around the PETH peak (Fig. 2B).

### Simulation of optogenetically evoked response latencies

To assess whether the suppression latency of T1-DANs evoked by optogenetic BFCN stimulation was quantitatively consistent with a disynaptic (BFCN → BF-GABA → T1-DAN) rather than a trisynaptic pathway, we modeled the expected suppression latency as the sum of the component delays along each pathway using a Monte Carlo simulation (3 × 10⁵ samples; each delay drawn independently on each iteration from rectified Gaussian distributions). The onset latency of BFCN activation was drawn from the measured light-evoked first-spike latency distribution (4.75 ± 1.5 ms, mean ± standard deviation; Fig. S4D). The BFCN→BF-GABA recruitment delay was drawn from the nicotinic EPSP onset latency measured at striatal cholinergic-to-GABAergic synapses (3.6 ± 1.6 ms) (*137*). BF-GABA output then reached the VTA after long-range axonal conduction over the ∼5 mm HDB–VTA distance (*133*), with an assumed conduction velocity of 2.0 ± 1.4 m/s, based on reported conduction velocities of basal forebrain long-range GABAergic projection neurons (*138*, *139*). T1-DAN inhibition was simulated from the IPSC onset latency measured in acute slices (3.40 ± 0.83 ms; Fig. 3I). Because the EPSP and IPSC onset latencies approximate the presynaptic-spike-to-postsynaptic-potential delay rather than the full spike-to-effect delay, the same term (1.0 ± 0.5 ms) was added at each synapse to account for the time from postsynaptic potential onset to the postsynaptic effect (a spike at the intermediate relay, or a detectable suppression of firing at the dopaminergic neuron). For the trisynaptic pathway, an additional glutamatergic relay was simulated, conservatively modeled with the same EPSP onset latency of 3.6 ± 1.6 ms.

### Trial-by-trial comparison of single unit responses

Trial-by-trial single unit cue responses were determined as the spike counts within the response window defined as the peak time ± peak width (full width at half maximum), calculated from the PETH aligned to the fixed go cue, and normalized by the neuron’s mean fixed go cue response. To assess the predictive value of neuronal responses for behavioral decisions, we fitted a generalized linear model with a binomial distribution (logistic regression, Fig. 4B and S11B), using normalized trial-by-trial new cue responses as predictors and behavioral choice as the dependent variable. Model predictions were used to compute receiver operating characteristic (ROC) curves, and discriminability was quantified as the area under the curve (AUC), estimated by 5-fold cross-validation: within each fold the model was fitted on the training trials and used to predict the held-out trials, and a single AUC was computed from the pooled out-of-fold predictions. To ensure that the superior performance of the combined BFCN + DAN predictor (Fig. S11B) did not simply reflect the use of two neurons rather than one, combined predictions from cross-type pairs (BFCN + DAN) were compared with those from within-type pairs of two simultaneously recorded neurons of the same type (two BFCNs, two T1-DANs, or two T2-DANs, Fig. 4B).

To test temporal relationships, we also fitted a time-shifted linear model (Fig. 4D) in which trends in behavioral responsiveness were modeled as a linear transform of the temporally shifted version of the neuronal responses (both smoothed by a Gaussian kernel, standard deviation of 1.4 trials, see Fig. S11D). The same time-shifted linear model was additionally fitted between simultaneously recorded pairs of neuronal response traces to quantify temporal relationships across neurons. Integer temporal shifts within the ± 10 trials range were evaluated exhaustively. For each shift, gain and bias parameters were estimated by least-squares optimization, and the shift yielding the minimal sum of squared residuals was selected. Model performance was quantified using the coefficient of determination (R²).

Trial-by-trial new-cue responses of BFCNs and DANs and the animal’s behavioral responsiveness were correlated across new-cue trials using Spearman correlation. Partial Spearman correlations were computed for simultaneously recorded pairs for each pair of these three variables while controlling for the remaining one (Fig. 4C and S11C). Noise correlation between simultaneously recorded BFCNs and DANs was assessed as the Spearman correlation between the trial-by-trial fixed cue responses relative to the average fixed cue responses of the given neurons (Fig. 2F).

### Joint peri-event time histogram

Joint peri-event time histograms (JPETHs) between simultaneously recorded BFCNs and DANs were computed aligned to cues, rewards or punishments (Fig. S8E-G), following (*140*). Trial-by-trial spike rasters binned at 1 ms time resolution were used to compute a joint spike count matrix, in which each element represented the number of trials with a BFCN spike at time t_y_ and a DAN spike at time t_x_. Joint probabilities were acquired by normalizing with the number of trials. To dissociate genuine temporal interactions from correlations driven by individual firing rate modulations, the expected joint activity under independence was estimated as the outer product of the marginal PETHs of the two neurons. The interaction matrix was then computed by subtracting this expectation from the observed joint probability matrix and normalizing by the product of the trial-by-trial standard deviations of the two spike trains, yielding a z-score–type interaction measure. The lag-dependence of BFCN-DAN interactions was quantified by averaging along the diagonals of the interaction matrix corresponding to fixed time lags between the two neurons. Finally, JPETHs and interactions were averaged in 20 ms non-overlapping bins for visualization and population analyses (Fig. 2C and S8E-G). For descriptive quantification, the population-average of these interaction profiles (Fig. 2D) was used to locate the lag ranges corresponding to the most prominent troughs and peaks (half-height width), and the average interaction within this range per pair was tested against no interaction (Fig. S8H; two-sided Wilcoxon signed-rank test).

### Cross-correlation

Cross-correlograms (CCGs) were computed for simultaneously recorded neuron pairs at 1 ms resolution and normalized by a Poisson baseline expectation. CCGs were computed either using all spikes (Fig. 2A) or using inter-trial intervals, to reveal potential network connections by eliminating the co-modulatory effects of behavioral events (Fig. 3A, S8A and S9). Normalized CCGs were smoothed by a Gaussian kernel (standard deviation, 15 ms).

For the photometry recordings, concurrently recorded continuous ACh and DA ΔF/F signals were cross-correlated following median filtering (100 ms window). The median cross-correlations across sessions were computed for each animal with simultaneous recordings (n = 11, including 5 control mice with simultaneous recordings from the chemogenetic cohort), normalized to the maximum value, and averaged across animals (Fig. 2E).

### Analysis of in vitro electrophysiology recordings

IPSC amplitude was measured as the baseline-subtracted peak current within a post-stimulus window. Latency was defined as the time from stimulus onset to 10% of peak amplitude and rise time as the time between 10–90% of peak amplitude. Paired-pulse ratio was calculated as pulse 2/pulse 1. TH+ and TH− groups were compared using two-sided Mann–Whitney U tests. Spike timing was detected using threshold-based methods and visualized with raster plots aligned to optical stimulus onset. Spike probability heatmaps were generated by aligning spikes to stimulus onset, binning spike times into 50 ms bins, normalizing counts to the maximum, and applying Gaussian smoothing (σ = 0.8) prior to heatmap visualization.

### Statistics

Statistical comparisons were performed using two-sided paired (Wilcoxon signed rank test) and two-sided non-paired (Mann-Whitney U test) non-parametric tests as appropriate, since normality of the data cannot be determined statistically. The SALT test (*16*) based on the Jensen-Shannon information divergence measure was used to assess significance of optogenetic tagging (p < 0.01). Box-whisker plots show median, interquartile range and non-outlier range. Individual data points were plotted wherever possible. Reliability of mean estimates is indicated by the standard error of the mean (SEM).

## Author contribution

B.H., B.K. and D.S. designed the experiments. B.K. established the behavioral task. V.P. and B.K. performed electrophysiology and fiber photometry experiments. Í.S. performed chemogenetic experiments. D.S. performed in vitro electrophysiology experiments. P.H. performed anterograde tracing. B.K. performed in vivo track reconstructions supervised by K.S.. Y.L. provided neurotransmitter biosensors. B.K. analyzed the data and prepared the figures. B.H. and B.K. wrote the manuscript with input from D.S. and Í.S. and Y.L.. B.H. supervised the project and acquired funding.

## Acknowledgements

We thank Katalin Lengyel, Ildikó Horváth, Annamária Benke and Weihao Sheng for technical assistance and Katalin Sviatkó, Sergio Martínez-Bellver, Nicola Solari and Diána Balázsfi for technical training. We thank Armin Lak for providing feedback on an earlier version of the manuscript and the FENS-Kavli Network of Excellence for fruitful discussions. We thank Norbert Hájos and Kinga Müller for providing viral constructs. We acknowledge the Nikon Center of Excellence at the HUN-REN Institute of Experimental Medicine (IEM), Nikon Europe, Nikon Austria, and Auro-Science Consulting for kindly providing microscopy support and the supportive help of the Central Virus Laboratory and the Behavioral Unit of IEM. This work was supported by the NKFIH K147097 grant of the National Research, Development and Innovation Office, the Hungarian Brain Research Program NAP3.0 (NAP2022-I-1/2022) and the “Lendület” LP2024-8/2024 grants by the Hungarian Academy of Sciences, and the European Research Council Starting Grant no. 715043 to B.H.; the ÚNKP-19-3, ÚNKP-20-3 and ÚNKP 21-3 New National Excellence Program of the Ministry for Innovation and Technology from the source of the National Research, Development and Innovation Fund and the Postdoctoral Fellowship of the European Molecular Biology Organization (EMBO, grant no. ALTF 699-2025) to B.K.; and the Postdoctoral Fellowship (R483-2024-1854) from the Lundbeck Foundation to D.S.

## Data availability

Data will be made available upon publication.

## References

1. W. Schultz, P. Dayan, P. R. Montague, A neural substrate of prediction and reward. Science 275, 1593–1599 (1997).

2. A. J. Yu, P. Dayan, Uncertainty, neuromodulation, and attention. Neuron 46 (2005).

3. K. Doya, Metalearning and neuromodulation. Neural Networks 15 (2002).

4. P. Hegedüs, K. Sviatkó, B. Király, S. Martínez-Bellver, B. Hangya, Cholinergic activity reflects reward expectations and predicts behavioral responses. iScience 26 (2023).

5. J. Sturgill, P. Hegedus, S. Li, Q. Chevy, A. Siebels, M. Jing, Y. Li, B. Hangya, A. Kepecs, Basal forebrain-derived acetylcholine encodes valence-free reinforcement prediction error. bioRxiv (2020).

6. V. Minces, L. Pinto, Y. Dan, A. A. Chiba, Cholinergic shaping of neural correlations. Proc. Natl. Acad. Sci. U. S. A. 114, 5725–5730 (2017).

7. R. B. Crouse, K. Kim, H. M. Batchelor, E. M. Girardi, R. Kamaletdinova, J. Chan, P. Rajebhosale, S. T. Pittenger, L. W. Role, D. A. Talmage, M. Jing, Y. Li, X. B. Gao, Y. S. Mineur, M. R. Picciotto, Acetylcholine is released in the basolateral amygdala in response to predictors of reward and enhances the learning of cue-reward contingency. Elife 9 (2020).

8. M. R. Roesch, G. R. Esber, J. Li, N. D. Daw, G. Schoenbaum, Surprise! Neural correlates of Pearce-Hall and Rescorla-Wagner coexist within the brain. Eur. J. Neurosci. 35 (2012).

9. J. J. Day, M. F. Roitman, R. M. Wightman, R. M. Carelli, Associative learning mediates dynamic shifts in dopamine signaling in the nucleus accumbens. Nat. Neurosci. 10 (2007).

10. I. Tsutsui-Kimura, H. Matsumoto, K. Akiti, M. M. Yamada, N. Uchida, M. Watabe-Uchida, Distinct temporal difference error signals in dopamine axons in three regions of the striatum in a decision-making task. Elife 9, 1–39 (2020).

11. W. Menegas, B. M. Babayan, N. Uchida, M. Watabe-Uchida, Opposite initialization to novel cues in dopamine signaling in ventral and posterior striatum in mice. Elife 6 (2017).

12. B. Hangya, S. P. Ranade, M. Lorenc, A. Kepecs, Central Cholinergic Neurons Are Rapidly Recruited by Reinforcement Feedback. Cell 162, 1155–1168 (2015).

13. T. C. Harrison, L. Pinto, J. R. Brock, Y. Dan, Calcium imaging of basal forebrain activity during innate and learned behaviors. Front. Neural Circuits 10 (2016).

14. M. Lovett-Barron, P. Kaifosh, M. A. Kheirbek, N. Danielson, J. D. Zaremba, T. R. Reardon, G. F. Turi, R. Hen, B. V. Zemelman, A. Losonczy, Dendritic inhibition in the hippocampus supports fear learning. Science (80-.). 343 (2014).

15. T. Laszlovszky, D. Schlingloff, P. Hegedüs, T. F. Freund, A. Gulyás, A. Kepecs, B. Hangya, Distinct synchronization, cortical coupling and behavioral function of two basal forebrain cholinergic neuron types. Nat. Neurosci. 23 (2020).

16. D. Kvitsiani, S. Ranade, B. Hangya, H. Taniguchi, J. Z. Huang, A. Kepecs, Distinct behavioural and network correlates of two interneuron types in prefrontal cortex. Nature 498, 363–6 (2013).

17. S. Q. Lima, T. Hromádka, P. Znamenskiy, A. M. Zador, PINP: a new method of tagging neuronal populations for identification during in vivo electrophysiological recording. PLoS One 4, e6099 (2009).

18. A. Széll, S. Martínez-Bellver, P. Hegedüs, B. Hangya, OPETH: Open Source Solution for Real-Time Peri-Event Time Histogram Based on Open Ephys. Front. Neuroinform. 14 (2020).

19. L. J. Agostinelli, J. C. Geerling, T. E. Scammell, Basal forebrain subcortical projections. Brain Struct. Funct. 224 (2019).

20. L. Zaborszky, A. van den Pol, E. Gyengesi, “The Basal Forebrain Cholinergic Projection System in Mice” in The Mouse Nervous System (Elsevier, 2012; https://linkinghub.elsevier.com/retrieve/pii/B9780123694973100287), pp. 684–718.

21. A. Aransay, C. Rodríguez-López, M. García-Amado, F. Clascá, L. Prensa, Long-range projection neurons of the mouse ventral tegmental area: A single-cell axon tracing analysis. Front. Neuroanat. 9 (2015).

22. B. Király, B. Hangya, Navigating the Statistical Minefield of Model Selection and Clustering in Neuroscience. eNeuro 9 (2022).

23. B. F. Sadacca, J. L. Jones, G. Schoenbaum, Midbrain dopamine neurons compute inferred and cached value prediction errors in a common framework. Elife 5 (2016).

24. J. Y. Cohen, S. Haesler, L. Vong, B. B. Lowell, N. Uchida, Neuron-type-specific signals for reward and punishment in the ventral tegmental area. (2012). 10.1038/nature10754.

25. M. Matsumoto, O. Hikosaka, Two types of dopamine neuron distinctly convey positive and negative motivational signals. Nature 459 (2009).

26. C. D. Fiorillo, Two dimensions of value: Dopamine neurons represent reward but not aversiveness. Science (80-.). 341 (2013).

27. M. Azcorra, Z. Gaertner, C. Davidson, Q. He, H. Kim, S. Nagappan, C. K. Hayes, C. Ramakrishnan, L. Fenno, Y. S. Kim, K. Deisseroth, R. Longnecker, R. Awatramani, D. A. Dombeck, Unique functional responses differentially map onto genetic subtypes of dopamine neurons. Nat. Neurosci. 26 (2023).

28. S. Lammel, B. K. Lim, C. Ran, K. W. Huang, M. J. Betley, K. M. Tye, K. Deisseroth, R. C. Malenka, Input-specific control of reward and aversion in the ventral tegmental area. Nature 491 (2012).

29. J. W. de Jong, Y. Liang, J. P. H. Verharen, K. M. Fraser, S. Lammel, State and rate-of-change encoding in parallel mesoaccumbal dopamine pathways. Nat. Neurosci. 27 (2024).

30. F. Brischoux, S. Chakraborty, D. I. Brierley, M. A. Ungless, Phasic excitation of dopamine neurons in ventral VTA by noxious stimuli. Proc. Natl. Acad. Sci. U. S. A. 106 (2009).

31. L. X. Cai, K. Pizano, G. W. Gundersen, C. L. Hayes, W. T. Fleming, S. Holt, J. M. Cox, I. B. Witten, Distinct signals in medial and lateral VTA dopamine neurons modulate fear extinction at different times. Elife 9 (2020).

32. R. S. Sutton, A. G. Barto, Reinforcement Learning: An Introduction. Cambridge: MIT Press. MA MIT Press. Sch. (1998).

33. Y. Niv, M. O. Duff, P. D. Dayan, Dopamine, uncertainty and TD learning. Behav. Brain Funct. 1 (2005).

34. R. N. Gentry, B. Lee, M. R. Roesch, Phasic dopamine release in the rat nucleus accumbens predicts approach and avoidance performance. Nat. Commun. 7 (2016).

35. E. B. Oleson, R. N. Gentry, V. C. Chioma, J. F. Cheer, Subsecond dopamine release in the nucleus accumbens predicts conditioned punishment and its successful avoidance. J. Neurosci. 32 (2012).

36. C. C. D. Shute, P. R. Lewis, Cholinesterase-containing systems of the brain of the rat. Nature 199 (1963).

37. H. Kimura, P. L. McGeer, J. H. Peng, E. G. McGeer, The central cholinergic system studied by choline acetyltransferase immunohistochemistry in the cat. J. Comp. Neurol. 200 (1981).

38. J. Lehmann, J. I. Nagy, S. Atmadja, H. C. Fibiger, The nucleus basalis magnocellularis: The origin of a cholinergic projection to the neocortex of the rat. Neuroscience 5 (1980).

39. C. B. Saper, Organization of cerebral cortical afferent systems in the rat. II. Magnocellular basal nucleus. J. Comp. Neurol. 222 (1984).

40. D. Corbett, R. A. Wise, Intracranial self-stimulation in relation to the ascending dopaminergic systems of the midbrain: A moveable electrode mapping study. Brain Res. 185 (1980).

41. W. Schultz, P. Apicella, T. Ljungberg, Responses of monkey dopamine neurons to reward and conditioned stimuli during successive steps of learning a delayed response task. J. Neurosci. 13 (1993).

42. B. J. Everitt, T. W. Robbins, Central cholinergic systems and cognition. Annu. Rev. Psychol. 48 (1997).

43. M. E. Hasselmo, M. Sarter, Modes and models of forebrain cholinergic neuromodulation of cognition. (2011). 10.1038/npp.2010.104.

44. V. Parikh, M. Sarter, “Cholinergic mediation of attention: Contributions of phasic and tonic increases in prefrontal cholinergic activity” in Annals of the New York Academy of Sciences (2008)vol. 1129.

45. A. A. Disney, C. Aoki, M. J. Hawken, Gain Modulation by Nicotine in Macaque V1. Neuron 56 (2007).

46. A. Nelson, R. Mooney, The Basal Forebrain and Motor Cortex Provide Convergent yet Distinct Movement-Related Inputs to the Auditory Cortex. Neuron 90 (2016).

47. L. Pinto, M. J. Goard, D. Estandian, M. Xu, A. C. Kwan, S. H. Lee, T. C. Harrison, G. Feng, Y. Dan, Fast modulation of visual perception by basal forebrain cholinergic neurons. Nat. Neurosci. 16 (2013).

48. S. Lohani, A. H. Moberly, H. Benisty, B. Landa, M. Jing, Y. Li, M. J. Higley, J. A. Cardin, Spatiotemporally heterogeneous coordination of cholinergic and neocortical activity. Nat. Neurosci. 25 (2022).

49. J. McGaughy, R. A. Koene, H. Eichenbaum, M. E. Hasselmo, Cholinergic deafferentation of the entorhinal cortex in rats impairs encoding of novel but not familiar stimuli in a delayed nonmatch-to-sample task. J. Neurosci. 25 (2005).

50. M. V. Puig, E. G. Antzoulatos, E. K. Miller, Prefrontal dopamine in associative learning and memory. (2014). 10.1016/j.neuroscience.2014.09.026.

51. M. G. Baxter, D. J. Bucci, L. K. Gorman, R. G. Wiley, M. Gallagher, Selective immunotoxic lesions of basal forebrain cholinergic cells: Effects on learning and memory in rats. Behav. Neurosci. 127 (2013).

52. L. S. Zweifel, J. G. Parker, C. J. Lobb, A. Rainwater, V. Wall, J. P. Fadok, M. Darvas, M. J. Kim, S. J. Y. Mizumori, C. A. Paladini, P. E. M. Phillips, R. D. Palmiter, Disruption of NMDAR-dependent burst firing by dopamine neurons provides selective assessment of phasic dopamine-dependent behavior. Proc. Natl. Acad. Sci. U. S. A. 106 (2009).

53. Z. Gu, J. L. Yakel, Timing-Dependent Septal Cholinergic Induction of Dynamic Hippocampal Synaptic Plasticity. Neuron 71 (2011).

54. J. L. Yakel, Nicotinic ACh receptors in the hippocampus: Role in excitability and plasticity. (2012). 10.1093/ntr/nts091.

55. A. Kirkwood, C. Rozas, J. Kirkwood, F. Perez, M. F. Bear, Modulation of long-term synaptic depression in visual cortex by acetylcholine and norepinephrine. J. Neurosci. 19 (1999).

56. R. C. Froemke, M. M. Merzenich, C. E. Schreiner, A synaptic memory trace for cortical receptive field plasticity. Nature 450 (2007).

57. M. W. Shiflett, B. W. Balleine, Molecular substrates of action control in cortico-striatal circuits. (2011). 10.1016/j.pneurobio.2011.05.007.

58. J. Urban-Ciecko, J. S. Jouhanneau, S. E. Myal, J. F. A. Poulet, A. L. Barth, Precisely Timed Nicotinic Activation Drives SST Inhibition in Neocortical Circuits. Neuron 97 (2018).

59. J. J. Letzkus, S. B. E. Wolff, E. M. M. Meyer, P. Tovote, J. Courtin, C. Herry, A. Lüthi, A disinhibitory microcircuit for associative fear learning in the auditory cortex. Nature 480 (2011).

60. J. Palacios-Filardo, J. R. Mellor, Neuromodulation of hippocampal long-term synaptic plasticity. (2019). 10.1016/j.conb.2018.08.009.

61. S. Zannone, Z. Brzosko, O. Paulsen, C. Clopath, Acetylcholine-modulated plasticity in reward-driven navigation: a computational study. Sci. Rep. 8 (2018).

62. F. Zhu, S. Elnozahy, J. Lawlor, K. V. Kuchibhotla, The cholinergic basal forebrain provides a parallel channel for state-dependent sensory signaling to auditory cortex. Nat. Neurosci. 26 (2023).

63. J. Y. Cohen, M. W. Amoroso, N. Uchida, Serotonergic neurons signal reward and punishment on multiple timescales. Elife, doi: 10.7554/eLife.06346 (2015).

64. D. Tse, R. F. Langston, M. Kakeyama, I. Bethus, P. A. Spooner, E. R. Wood, M. P. Witter, R. G. M. Morris, Schemas and memory consolidation. Science (80-.). 316 (2007).

65. B. Király, D. Balázsfi, I. Horváth, N. Solari, K. Sviatkó, K. Lengyel, E. Birtalan, M. Babos, G. Bagaméry, D. Máthé, K. Szigeti, B. Hangya, In vivo localization of chronically implanted electrodes and optic fibers in mice. Nat. Commun. 11, 4686 (2020).

66. M. R. Gielow, L. Zaborszky, The Input-Output Relationship of the Cholinergic Basal Forebrain. Cell Rep. 18 (2017).

67. I. Villano, A. Messina, A. Valenzano, F. Moscatelli, T. Esposito, V. Monda, M. Esposito, F. Precenzano, M. Carotenuto, A. Viggiano, S. Chieffi, G. Cibelli, M. Monda, G. Messina, Basal forebrain cholinergic system and orexin neurons: Effects on attention. (2017). 10.3389/fnbeh.2017.00010.

68. J. Tian, R. Huang, J. Y. Cohen, F. Osakada, D. Kobak, C. K. Machens, E. M. Callaway, N. Uchida, M. Watabe-Uchida, Distributed and Mixed Information in Monosynaptic Inputs to Dopamine Neurons. Neuron 91 (2016).

69. M. Watabe-Uchida, L. Zhu, S. K. Ogawa, A. Vamanrao, N. Uchida, Whole-Brain Mapping of Direct Inputs to Midbrain Dopamine Neurons. Neuron 74 (2012).

70. W. Menegas, J. F. Bergan, S. K. Ogawa, Y. Isogai, K. U. Venkataraju, P. Osten, N. Uchida, M. Watabe-Uchida, Dopamine neurons projecting to the posterior striatum form an anatomically distinct subclass. Elife 4, 1–30 (2015).

71. L. Zaborszky, A. Csordas, K. Mosca, J. Kim, M. R. Gielow, C. Vadasz, Z. Nadasdy, Neurons in the basal forebrain project to the cortex in a complex topographic organization that reflects corticocortical connectivity patterns: An experimental study based on retrograde tracing and 3D reconstruction. Cereb. Cortex 25 (2015).

72. Y. Hu, J. Zylberberg, E. Shea-Brown, The Sign Rule and Beyond: Boundary Effects, Flexibility, and Noise Correlations in Neural Population Codes. PLoS Comput. Biol. 10 (2014).

73. I. Kanitscheider, R. Coen-Cagli, A. Pouget, Origin of information-limiting noise correlations. Proc. Natl. Acad. Sci. U. S. A. 112 (2015).

74. D. Dautan, A. S. Souza, I. Huerta-Ocampo, M. Valencia, M. Assous, I. B. Witten, K. Deisseroth, J. M. Tepper, J. P. Bolam, T. V. Gerdjikov, J. Mena-Segovia, Segregated cholinergic transmission modulates dopamine neurons integrated in distinct functional circuits. Nat. Neurosci. 19 (2016).

75. I. Avila, S. C. Lin, Motivational Salience Signal in the Basal Forebrain Is Coupled with Faster and More Precise Decision Speed. PLoS Biol. 12 (2014).

76. S. C. Lin, R. E. Brown, M. G. H. Shuler, C. C. H. Petersen, A. Kepecs, Optogenetic dissection of the basal forebrain neuromodulatory control of cortical activation, plasticity, and cognition. J. Neurosci. 35 (2015).

77. S. C. Lin, M. A. L. Nicolelis, Neuronal Ensemble Bursting in the Basal Forebrain Encodes Salience Irrespective of Valence. Neuron 59 (2008).

78. L. Faget, L. Oriol, W. C. Lee, V. Zell, C. Sargent, A. Flores, N. G. Hollon, D. Ramanathan, T. S. Hnasko, Ventral pallidum GABA and glutamate neurons drive approach and avoidance through distinct modulation of VTA cell types. Nat. Commun. 15 (2024).

79. S. Lammel, D. I. Ion, J. Roeper, R. C. Malenka, Projection-Specific Modulation of Dopamine Neuron Synapses by Aversive and Rewarding Stimuli. Neuron 70 (2011).

80. N. F. Parker, C. M. Cameron, J. P. Taliaferro, J. Lee, J. Y. Choi, T. J. Davidson, N. D. Daw, I. B. Witten, Reward and choice encoding in terminals of midbrain dopamine neurons depends on striatal target. Nat. Neurosci. 19 (2016).

81. W. Menegas, K. Akiti, R. Amo, N. Uchida, M. Watabe-Uchida, Dopamine neurons projecting to the posterior striatum reinforce avoidance of threatening stimuli. Nat. Neurosci. 21, 1421–1430 (2018).

82. S.-J. Bouchard, J. Bouchard, M. Levesque, V. Breton-Provencher, System-wide dissociation of reward and aversive dopaminergic signals. bioRxiv (2025).

83. H. Kim, C. Xu, C. Washington, C. Shi, M. Lowman, J. M. Kebschull, POINTseq: Cell-type-specific barcoding reveals single-cell projection architecture of the mouse dopaminergic system. Neuron, doi: 10.1016/j.neuron.2026.05.005 (2026).

84. M. A. Klitenick, A. Y. Deutch, L. Churchill, P. W. Kalivas, Topography and functional role of dopaminergic projections from the ventral mesencephalic tegmentum to the ventral pallidum. Neuroscience 50 (1992).

85. Y.-J. Jhong, M.-C. Chiang, A. Guillaumin, C.-Y. Cheng, M.-E. Kuo, C.-Z. Lee, A. Scaglione, F. Georges, S.-C. Lin, Dopamine neurons drive basal forebrain reward prediction in milliseconds. bioRxiv, 2026.10.01.755874 (2026).

86. S. Threlfell, T. Lalic, N. J. Platt, K. A. Jennings, K. Deisseroth, S. J. Cragg, Striatal dopamine release is triggered by synchronized activity in cholinergic interneurons. Neuron 75 (2012).

87. A. Mohebi, V. L. Collins, J. D. Berke, Accumbens cholinergic interneurons dynamically promote dopamine release and enable motivation. Elife 12 (2023).

88. G. C. Touponse, M. B. Pomrenze, T. Yassine, V. Mehta, N. Denomme, Z. Zhang, R. C. Malenka, N. Eshel, Cholinergic modulation of dopamine release drives effortful behavior. bioRxiv (2025).

89. L. Chantranupong, C. C. Beron, J. A. Zimmer, M. J. Wen, W. Wang, B. L. Sabatini, Dopamine and glutamate regulate striatal acetylcholine in decision-making. Nature 621 (2023).

90. L. Matityahu, N. Gilin, G. A. Sarpong, Y. Atamna, L. Tiroshi, N. X. Tritsch, J. R. Wickens, J. A. Goldberg, Acetylcholine waves and dopamine release in the striatum. Nat. Commun. 14 (2023).

91. S. Bouabid, L. Zhang, M. A. T. Vu, K. Tang, B. M. Graham, C. A. Noggle, M. W. Howe, Distinct spatially organized striatum-wide acetylcholine dynamics for the learning and extinction of Pavlovian associations. Nat. Commun. 16 (2025).

92. J. O’Keefe, M. L. Recce, Phase relationship between hippocampal place units and the EEG theta rhythm. Hippocampus 3, 317–330 (1993).

93. W. E. Skaggs, B. L. McNaughton, M. A. Wilson, C. A. Barnes, Theta phase precession in hippocampal neuronal populations and the compression of temporal sequences. Hippocampus 6 (1996).

94. A. Johnson, A. D. Redish, Neural ensembles in CA3 transiently encode paths forward of the animal at a decision point. J. Neurosci. 27 (2007).

95. G. Buzsáki, D. Tingley, Space and Time: The Hippocampus as a Sequence Generator. (2018). 10.1016/j.tics.2018.07.006.

96. W. Guo, B. Robert, D. B. Polley, The Cholinergic Basal Forebrain Links Auditory Stimuli with Delayed Reinforcement to Support Learning. Neuron 103 (2019).

97. A. Atri, K. A. Norman, M. M. Nicolas, S. C. Cramer, M. E. Hasselmo, S. Sherman, B. A. Kirchhoff, M. D. Greicius, H. C. Breiter, C. E. Stern, Blockade of Central Cholinergic Receptors Impairs New Learning and Increases Proactive Interference in a Word Paired-Associate Memory Task. Behav. Neurosci. 118 (2004).

98. T. Gedankien, J. Kriegel, E. Zabeh, D. McDonagh, B. Lega, J. Jacobs, Cholinergic blockade reveals role for human hippocampal theta in encoding but not retrieval. (2025).

99. W. Schultz, Predictive reward signal of dopamine neurons. (1998). 10.1152/jn.1998.80.1.1.

100. R. A. Wise, Dopamine, learning and motivation. (2004). 10.1038/nrn1406.

101. Z. Su, J. Y. Cohen, Two types of locus coeruleus norepinephrine neurons drive reinforcement learning. bioRxiv (2022).

102. P. J. Whitehouse, D. L. Price, R. G. Struble, A. W. Clark, J. T. Coyle, M. R. DeLong, Alzheimer’s disease and senile dementia: Loss of neurons in the basal forebrain. Science (80-.). 215 (1982).

103. M. Mesulam, The Cholinergic Lesion of Alzheimer’s Disease: Pivotal Factor or Side Show? (2004). 10.1101/lm.69204.

104. M. M. Mesulam, Alzheimer plaques and cortical cholinergic innervation. Neuroscience 17 (1986).

105. I. Iraizoz, S. de Lacalle, L. M. Gonzalo, Cell loss and nuclear hypertrophy in topographical subdivisions of the nucleus basalis of Meynert in Alzheimer’s disease. Neuroscience 41 (1991).

106. M. Gallagher, P. J. Colombo, Ageing: the cholinergic hypothesis of cognitive decline. Curr. Opin. Neurobiol. 5 (1995).

107. S. Allard, M. G. Hussain Shuler, Cholinergic reinforcement signaling is impaired by amyloidosis prior to its synaptic loss. J. Neurosci. 43 (2023).

108. H. Bernheimer, W. Birkmayer, O. Hornykiewicz, K. Jellinger, F. Seitelberger, Brain dopamine and the syndromes of Parkinson and Huntington Clinical, morphological and neurochemical correlations. J. Neurol. Sci. 20 (1973).

109. D. Lindholm, J. Mäkelä, V. Di Liberto, G. Mudò, N. Belluardo, O. Eriksson, M. Saarma, Current disease modifying approaches to treat Parkinson’s disease. (2016). 10.1007/s00018-015-2101-1.

110. K. R. Gamble, T. J. Cummings, S. E. Lo, P. T. Ghosh, J. H. Howard, D. V. Howard, Implicit sequence learning in people with Parkinson’s disease. Front. Hum. Neurosci. 8 (2014).

111. K. R. Chaudhuri, A. H. Schapira, Non-motor symptoms of Parkinson’s disease: dopaminergic pathophysiology and treatment. (2009). 10.1016/S1474-4422(09)70068-7.

112. A. A. Kehagia, R. A. Barker, T. W. Robbins, Neuropsychological and clinical heterogeneity of cognitive impairment and dementia in patients with Parkinson’s disease. (2010). 10.1016/S1474-4422(10)70212-X.

113. W. Dauer, S. Przedborski, Parkinson’s disease: Mechanisms and models. (2003). 10.1016/S0896-6273(03)00568-3.

114. M. J. Frank, L. C. Seeberger, R. C. O’Reilly, By carrot or by stick: Cognitive reinforcement learning in Parkinsonism. Science (80-.). 306 (2004).

115. A. Fushiki, D. Ng, Z. R. Lewis, A. Yadav, T. Saraiva, L. A. Hammand, C. Wirblich, B. Tasic, V. Menon, J. A. da Silva, R. M. Costa, A Vulnerable Subtype of Dopaminergic Neurons Drives Early Motor Deficits in Parkinson’s Disease. bioRxiv Prepr. Serv. Biol., doi: 10.1101/2024.12.20.629776 (2024).

116. G. Liu, J. Yu, J. Ding, C. Xie, L. Sun, I. Rudenko, W. Zheng, N. Sastry, J. Luo, G. Rudow, J. C. Troncoso, H. Cai, Aldehyde dehydrogenase 1 defines and protects a nigrostriatal dopaminergic neuron subpopulation. J. Clin. Invest. 124 (2014).

117. M. Pereira Luppi, M. Azcorra, G. Caronia-Brown, J. F. Poulin, Z. Gaertner, S. Gatica, O. A. Moreno-Ramos, N. Nouri, M. Dubois, Y. C. Ma, C. Ramakrishnan, L. Fenno, Y. S. Kim, K. Deisseroth, F. Cicchetti, D. A. Dombeck, R. Awatramani, Sox6 expression distinguishes dorsally and ventrally biased dopamine neurons in the substantia nigra with distinctive properties and embryonic origins. Cell Rep. 37 (2021).

118. N. I. Bohnen, P. Kanel, M. L. T. M. Müller, “Molecular Imaging of the Cholinergic System in Parkinson’s Disease” in International Review of Neurobiology (2018)vol. 141.

119. P. J. Whitehouse, J. C. Hedreen, C. L. White, D. L. Price, Basal forebrain neurons in the dementia of Parkinson disease. Ann. Neurol. 13 (1983).

120. M. J. Barrett, S. A. Sperling, J. C. Blair, C. S. Freeman, J. L. Flanigan, M. E. Smolkin, C. A. Manning, T. J. Druzgal, Lower volume, more impairment: Reduced cholinergic basal forebrain grey matter density is associated with impaired cognition in Parkinson disease. J. Neurol. Neurosurg. Psychiatry 90 (2019).

121. L. Batzu, D. Urso, M. J. Grothe, D. Veréb, K. R. Chaudhuri, J. B. Pereira, Increased basal forebrain volumes could prevent cognitive decline in LRRK2 Parkinson’s disease. Neurobiol. Dis. 183 (2023).

122. C. Legault-Denis, M. Aghourian, J. P. Soucy, P. Rosa-Neto, A. Dagher, E. Aumont, R. Wickens, M. A. Bedard, Normal cognition in Parkinson’s disease may involve hippocampal cholinergic compensation: An exploratory PET imaging study with [18F]-FEOBV. Park. Relat. Disord. 91 (2021).

123. N. I. Bohnen, A. J. Yarnall, R. S. Weil, E. Moro, M. S. Moehle, P. Borghammer, M. A. Bedard, R. L. Albin, Cholinergic system changes in Parkinson’s disease: emerging therapeutic approaches. (2022). 10.1016/S1474-4422(21)00377-X.

124. A. Nobili, E. C. Latagliata, M. T. Viscomi, V. Cavallucci, D. Cutuli, G. Giacovazzo, P. Krashia, F. R. Rizzo, R. Marino, M. Federici, P. De Bartolo, D. Aversa, M. C. Dell’Acqua, A. Cordella, M. Sancandi, F. Keller, L. Petrosini, S. Puglisi-Allegra, N. B. Mercuri, R. Coccurello, N. Berretta, M. D’Amelio, Dopamine neuronal loss contributes to memory and reward dysfunction in a model of Alzheimer’s disease. Nat. Commun. 8 (2017).

125. L. Serra, M. D’Amelio, S. Esposito, C. Di Domenico, G. Koch, C. Marra, N. B. Mercuri, C. Caltagirone, C. A. Artusi, L. Lopiano, M. Cercignani, M. Bozzali, Ventral tegmental area disconnection contributes two years early to correctly classify patients converted to alzheimer’s disease: Implications for treatment. J. Alzheimer’s Dis. 82 (2021).

126. K. Henjum, L. O. Watne, K. Godang, N. B. Halaas, R. S. Eldholm, K. Blennow, H. Zetterberg, I. Saltvedt, J. Bollerslev, A. B. Knapskog, Cerebrospinal fluid catecholamines in Alzheimer’s disease patients with and without biological disease. Transl. Psychiatry 12 (2022).

127. J. E. Ferguson, C. Boldt, A. D. Redish, Creating low-impedance tetrodes by electroplating with additives. Sensors Actuators, A Phys. 156 (2009).

128. N. Solari, K. Sviatkó, T. Laszlovszky, P. Hegedüs, B. Hangya, Open Source Tools for Temporally Controlled Rodent Behavior Suitable for Electrophysiology and Optogenetic Manipulations. Front. Syst. Neurosci. 12 (2018).

129. F. Najafi, A. Giovannucci, S. S. H. Wang, J. F. Medina, Coding of stimulus strength via analog calcium signals in Purkinje cell dendrites of awake mice. Elife 3 (2014).

130. R. S. Nickerson, D. W. Burnham, Response times with nonaging foreperiods. J. Exp. Psychol. 79 (1969).

131. R. Näätänen, Non-aging fore-periods and simple reaction time. Acta Psychol. (Amst). 35 (1971).

132. J. Bai, T. L. H. Trinh, K. H. Chuang, A. Qiu, Atlas-based automatic mouse brain image segmentation revisited: Model complexity vs. image registration. Magn. Reson. Imaging, doi: 10.1016/j.mri.2012.02.010 (2012).

133. G. Paxinos, K. B. J. Franklin, The Mouse Brain in Stereotaxic Coordinates, 2nd Edition (2001).

134. M. Jendryka, M. Palchaudhuri, D. Ursu, B. van der Veen, B. Liss, D. Kätzel, W. Nissen, A. Pekcec, Pharmacokinetic and pharmacodynamic actions of clozapine-N-oxide, clozapine, and compound 21 in DREADD-based chemogenetics in mice. Sci. Rep. 9 (2019).

135. T. N. Lerner, C. Shilyansky, T. J. Davidson, K. E. Evans, K. T. Beier, K. A. Zalocusky, A. K. Crow, R. C. Malenka, L. Luo, R. Tomer, K. Deisseroth, Intact-Brain Analyses Reveal Distinct Information Carried by SNc Dopamine Subcircuits. Cell 162 (2015).

136. N. Schmitzer-Torbert, J. Jackson, D. Henze, K. Harris, A. D. Redish, Quantitative measures of cluster quality for use in extracellular recordings. Neuroscience 131 (2005).

137. D. F. English, O. Ibanez-Sandoval, E. Stark, F. Tecuapetla, G. Buzsáki, K. Deisseroth, J. M. Tepper, T. Koos, GABAergic circuits mediate the reinforcement-related signals of striatal cholinergic interneurons. Nat. Neurosci. 15 (2012).

138. Y. Lamour, P. Dutar, O. Rascol, A. Jobert, Basal forebrain neurons projecting to the rat frontoparietal cortex: Electrophysiological and pharmacological properties. Brain Res. 362 (1986).

139. G. A. Jones, S. K. Norris, Z. Henderson, Conduction velocities and membrane properties of different classes of rat septohippocampal neurons recorded in vitro. J. Physiol. 517 (1999).

140. A. M. H. J. Aertsen, G. L. Gerstein, M. K. Habib, G. Palm, Dynamics of neuronal firing correlation: Modulation of “effective connectivity.” J. Neurophysiol. 61 (1989).

