## Supplementary Figures for "Cholinergic–dopaminergic synergy and antagonism shape prediction-error broadcasting"

*Király et al.*

**Supplementary figures**

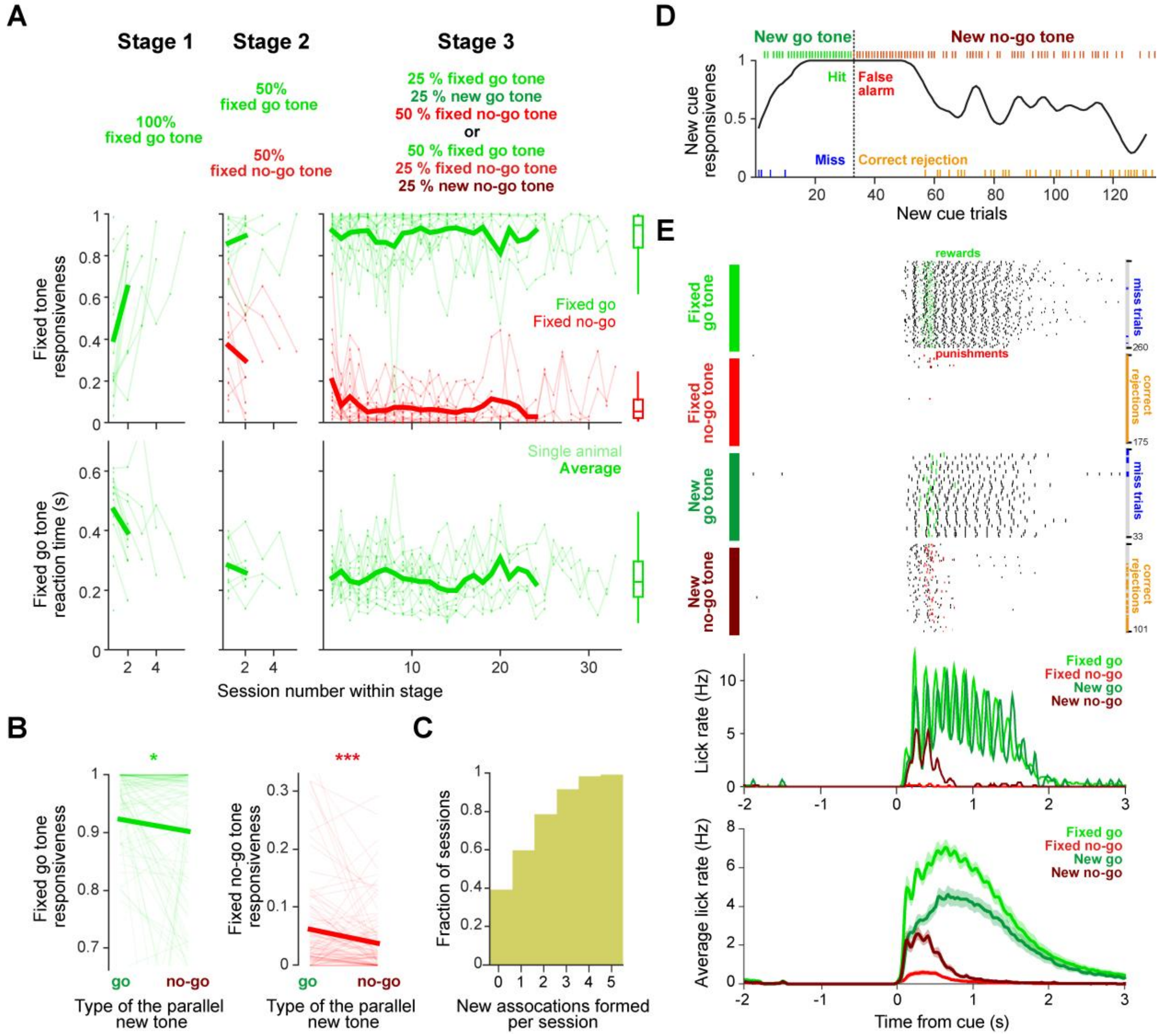


**Figure S1 Mice learn to form novel stimulus-action-outcome associations while they maintain stable performance on fixed associations.**

**A** Top, proportion of fixed and novel tones in different task phases. In Stage 1, mice are only introduced to the fixed go association. Once 70% performance is reached, mice proceed to Stage 2, in which the fixed no-go tone is introduced in 50% of the trials. After mice again reach 70% performance (>70% response rate to go tones and <30% response rate to no-go tones), mice enter Stage 3 (final stage), where half of either the fixed go or the fixed no-go tones are replaced with a new tone, keeping the overall number of go and no-go tones equal. Once 70% performance is reached, a new association replaces the former 'novel association', with randomized stimulus tone frequency and outcome (go or no-go). Middle, mice can learn the fixed go association (green) in 1-6 days (Stage 1) and the fixed no-go association (red) in an additional 1-5 days (Stage 2) and maintain stable fixed association performance throughout the final stage of the task. Bottom, reaction times decrease during the first two stages, after which they settle at a constant level (around 0.2 s for go tones). Thin lines, individual mice; thick lines, average (calculated from >5 individual mice). Box-whisker plots show median, interquartile range and non-outlier range across sessions of the final stage. **B** Responsiveness to both fixed tones slightly decreases, when the novel tone is a no-go tone compared to novel go tones (*, $p = 0.0164$; ***, $p = 0.00023$, two-sided paired Wilcoxon test). **C** Cumulative distribution function of the number of new associations formed per session. **D** Response rate to the novel tone (smoothed by a Gaussian kernel, standard deviation of 1.4 trials) from an example session, in which two new associations were learned (go then no-go, separated by the dashed line). Ticks indicate single trial decisions. Green, hit (response to the go tone); blue, missed go tone; red, false alarm (response to the no-go tone); orange, correct rejection of the no-go tone. **E** Top, lick rasters aligned to cue onset for the four association types in the same example session as in panel D. Corresponding changes in miss and correct-rejection trial density are shown on the right axis. Reward and punishment times in hit and false alarm trials are marked by green and red ticks on the corresponding raster plots. Note the increasing and decreasing reward anticipation as new go and no-go associations are formed. Middle, corresponding lick PETHs of the session. Bottom, grand-average PETHs across animals. Error shades represent SEM. Note the stable, high vs. near-zero reward anticipation to fixed go and no-go tones, respectively.

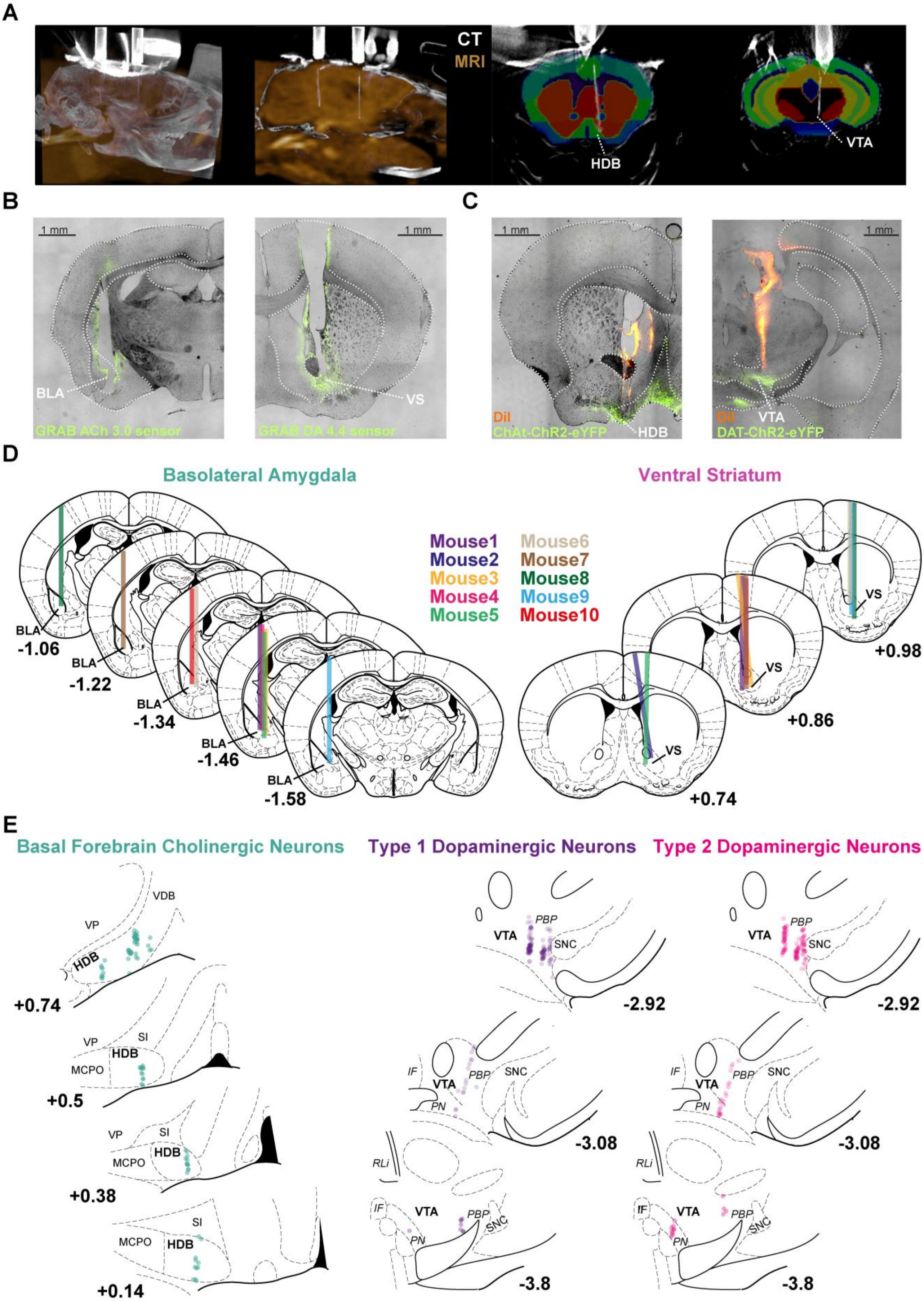
A
CT
MRI
HDB
VTA
B
C
1 mm
BLA
VS
GRAB ACh 3.0 sensor
GRAB DA 4.4 sensor
DiI
ChAt-ChR2-eYFP
HDB
VTA
DAT-ChR2-eYFP
D
Basolateral Amygdala
Ventral Striatum
Mouse1
Mouse2
Mouse3
Mouse4
Mouse5
Mouse6
Mouse7
Mouse8
Mouse9
Mouse10
BLA
-1.06
-1.22
-1.34
-1.46
-1.58
VS
+0.98
+0.86
+0.74
E
Basal Forebrain Cholinergic Neurons
Type 1 Dopaminergic Neurons
Type 2 Dopaminergic Neurons
VDB
VP
HDB
SI
MCPO
+0.74
+0.5
+0.38
+0.14
VTA
PBP
SNC
IF
PN
RLi
-2.92
-3.08
-3.8

**Figure S2 Reconstruction of implant positions.**

**A** In vivo localization of tetrode implants targeting the HDB and the VTA, based on preoperative MRI (gold) and postoperative CT (white) scans co-registered to a 3D mouse brain atlas (colored). Images from left to right, maximum intensity projection, sagittal slice, coronal slices. **B** Overlayed bright field and fluorescent micrographs of fiber photometry optic fiber tracks targeting the BLA (left) and the VS (right). Green, GRAB ACh3.0 or GRAB DA2m neuromodulator sensor expression. **C** Overlayed bright field and fluorescent micrographs of electrophysiological tetrode tracks targeting the HDB (left) and the VTA (right). Green, ChR2-eYFP expression. Red, DiI. **D** Reconstructed optic fiber tracks. Numbers reflect the anteroposterior coordinate from Bregma. **E** Reconstructed cholinergic and dopaminergic cell positions. Numbers reflect the anteroposterior coordinate from Bregma. HDB, horizontal limb of the diagonal band of Broca; MCPO, magnocellular preoptic nucleus; SI, substantia innominata; VDB, vertical limb of the diagonal band of Broca; VP, ventral pallidum; VTA, ventral tegmental area; IF, interfascicular nucleus; PN, paranigral nucleus; PBP, parabrachial pigmented nucleus; RLi, rostral linear nucleus; SNc, substantia nigra pars compacta; BLA, basolateral amygdala; VS, ventral striatum.

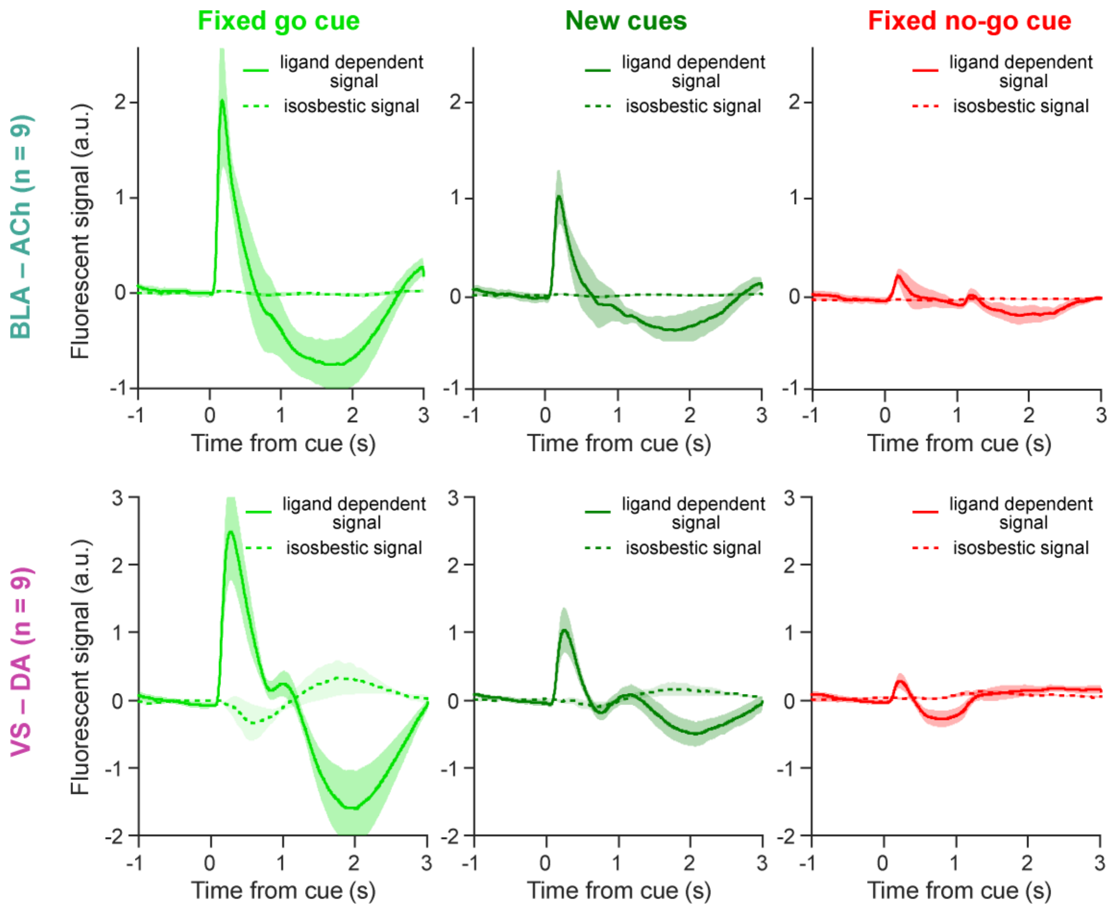


**Figure S3 Ligand-dependent and isosbestic fiber photometry signals.**

Top, cue-aligned PETHs of ACh signals from the basolateral amygdala averaged across animals (n = 9). Bottom, the same for DA signals from the ventral striatum (n = 9). Ligand-dependent signals, but not the isosbestic controls (dashed lines), scale with the reward-predictive value of the different tones, from the fixed go tone (left, largest), through new cues (middle, intermediate), to the fixed no-go tone (right, smallest). Error shades show SEM.

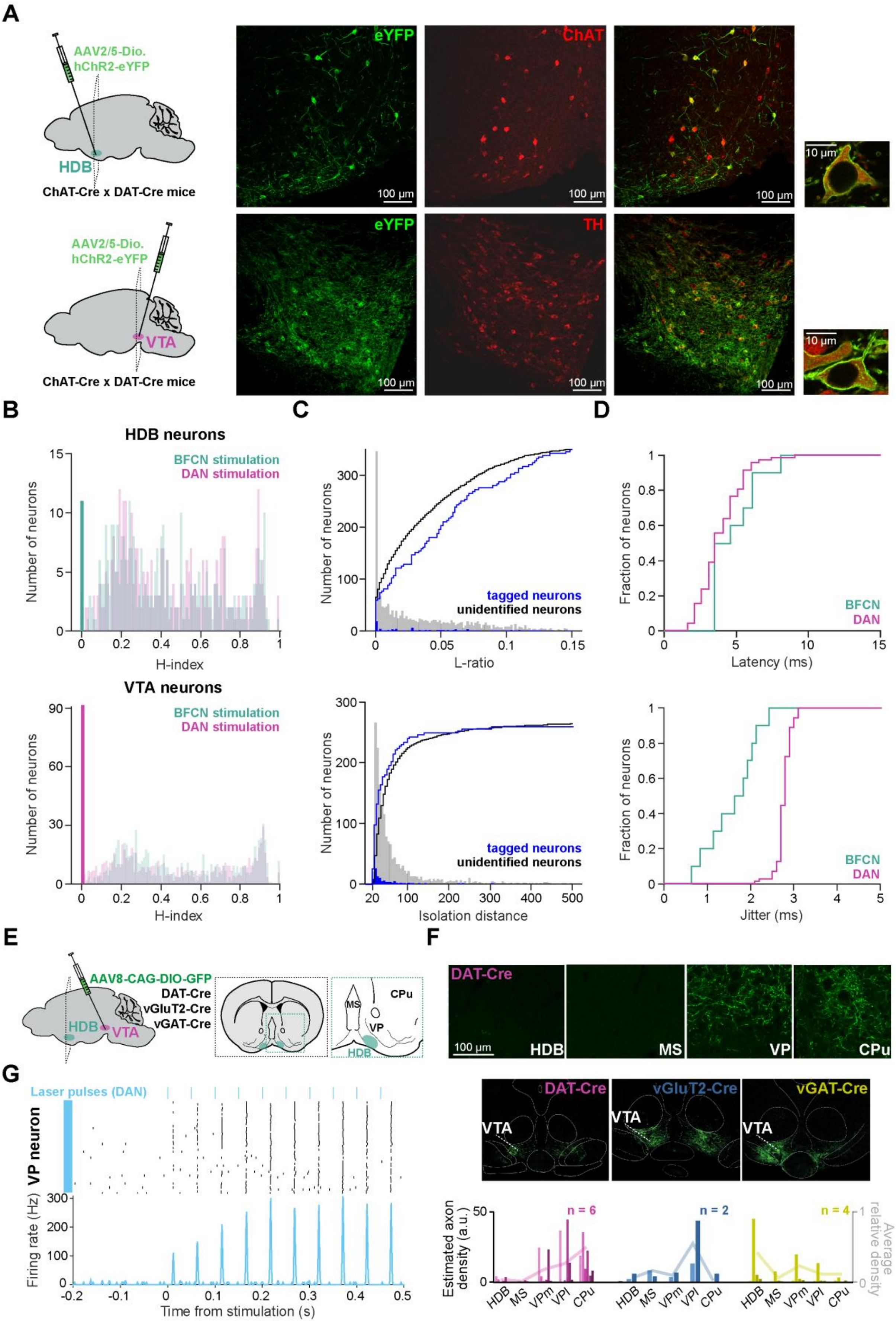
A
AAV2/5-Dio. hChR2-eYFP
HDB
ChAT-Cre x DAT-Cre mice
eYFP
ChAT
100 µm
10 µm
AAV2/5-Dio. hChR2-eYFP
VTA
ChAT-Cre x DAT-Cre mice
eYFP
TH
B
HDB neurons
BFCN stimulation
DAN stimulation
Number of neurons
H-index
VTA neurons
C
Number of neurons
tagged neurons
unidentified neurons
L-ratio
Isolation distance
D
Fraction of neurons
BFCN
DAN
Latency (ms)
Jitter (ms)
E
AAV8-CAG-DIO-GFP
DAT-Cre
vGluT2-Cre
vGAT-Cre
HDB
VTA
MS
CPu
VP
F
DAT-Cre
HDB
MS
VP
CPu
DAT-Cre
vGluT2-Cre
vGAT-Cre
VTA
Estimated axon density (a.u.)
Average relative density
n = 6
n = 2
n = 4
HDB
MS
VPm
VPl
CPu
G
Laser pulses (DAN)
VP neuron
Firing rate (Hz)
Time from stimulation (s)

**Figure S4 Optogenetic identification of cholinergic and dopaminergic neurons.**

**A** ChAT-Cre and DAT-Cre mice were crossed for optogenetic tagging of both cholinergic (top) and dopaminergic (bottom) neurons in the same animal. Fluorescent images show channelrhodopsin-expressing neurons (green, eYFP), ChAT-expressing cholinergic neurons in the HDB (top, ChAT-immunostaining in red) and tyrosine-hydroxylase- (TH-) expressing dopaminergic neurons in the VTA (bottom, TH-immunostaining in red). Overlayed images verify that channelrhodopsin-expressing neurons are ChAT+ in the HDB (top) and TH+ in the VTA (bottom). **B** Distribution of the significance values of the SALT test (stimulus-associated spike latency test, H-index) during stimulation of either brain areas (teal, HDB; magenta, VTA), for all HDB (top) and VTA (bottom) neurons recorded from mice with optogenetically identified neurons. The bright teal and bright magenta bars represent significantly light-responsive tagged cholinergic and dopaminergic neurons, respectively ($p < 0.01$). As expected based on the lack of direct neuromodulatory projections between HDB and VTA, the dual tagging approach did not lead to confounding effects. **C** Distribution of cluster quality measures L-ratio (top) and isolation distance (bottom) for tagged (blue) and non-tagged neurons (black). **D** Top, cumulative histogram of the response peak latency of cholinergic (teal) and dopaminergic (magenta) neurons after optogenetic stimulation. Bottom, cumulative histogram of the jitter of spike responses after optogenetic stimulation. **E** Schematic of the anterograde tracing experiments performed in DAT-Cre, vGluT2-Cre and vGAT-Cre animals to characterize dopaminergic, glutamatergic and GABAergic VTA projections to the basal forebrain. **F** Representative fluorescent images of dopaminergic axons in different brain areas. MS, medial septum; HDB, horizontal limb of the diagonal band of Broca; VP, ventral pallidum; CPu, caudate putamen. Middle, example injection sites in the VTA of DAT-Cre (left), vGluT2-Cre (middle) and vGAT-Cre (right) animals. Bottom, average relative density of VTA axons across the examined basal forebrain/basal ganglia regions (lines) in DAT-Cre (left), vGluT2-Cre (middle) and vGAT-Cre (right) animals. Bars show absolute axon density in individual mice (represented by different colors). VPm, medial ventral pallidum; VPl, lateral ventral pallidum. **G** Spike raster (top) and PETH (bottom) of an indirectly optogenetically activated example VP neuron aligned to VTA-DAN stimulation (blue lines).

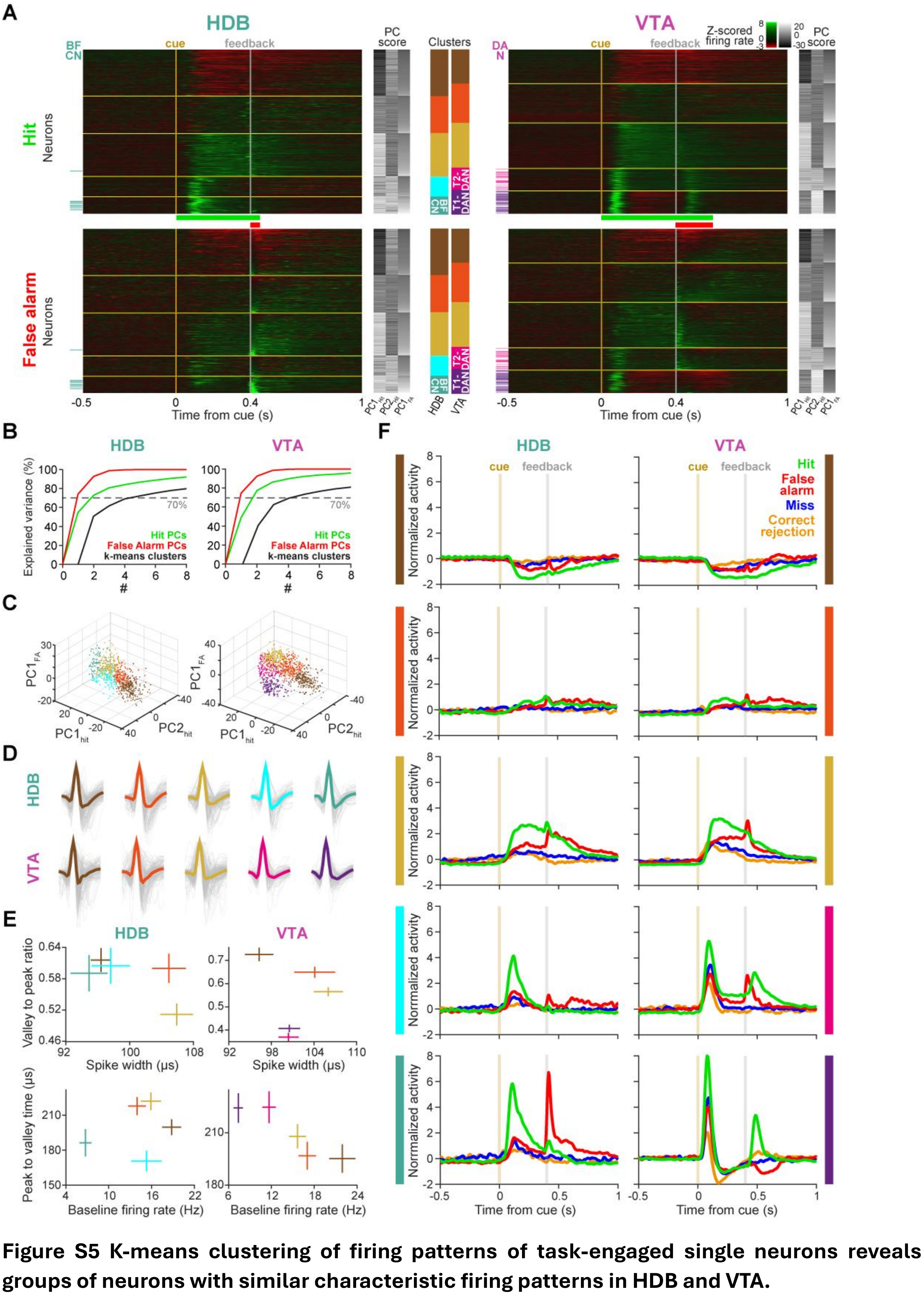

**Figure S5 K-means clustering of firing patterns of task-engaged single neurons reveals groups of neurons with similar characteristic firing patterns in HDB and VTA.**

**A** Z-scored PETHs of all HDB (left) and VTA (right) single units in hit (top) and false alarm trials (bottom). Green and red, firing rate above and below baseline, respectively. From -0.5 s to 0.4 s, PETHs are aligned to cue onset (0 s, mustard line). From 0.4 s to 1 s, PETHs are aligned to feedback time (0.4 s, gray line). Principal component analysis (PCA) was used for dimensionality reduction of the time series data. Greyscale heat maps on the right show principal component scores. K-means clustering was performed in the space spanned by the first two principal components of the hit PETHs and the first principal component of the false alarm PETHs, yielding five distinct clusters. Neurons were sorted according to their clusters. Within each cluster, neurons were sorted by the first principal component of the false alarm PETHs. Horizontal bars in the middle show the time windows used for clustering in hit (green) and false alarm (red) trials, chosen to capture the time scale of cholinergic and dopaminergic responses. Colored ticks on the left mark optogenetically identified cholinergic and dopaminergic neurons, almost exclusively falling into the last HDB (n = 10/11) and the last two VTA clusters (n = 79/79), corresponding to putative cholinergic and dopaminergic clusters, respectively. Vertical colored stacked bars in the middle indicate the proportion of neurons in each response cluster. **B** The elbow method was applied to the explained variance of the first 8 principal components and the number of clusters, showing that the majority (>70%) of the variance was explained by the first two principal components of the hit PETHs (green), the first principal component of the false alarm PETHs (red) and five clusters in both areas (left, HDB; right, VTA). **C** Scatter plot of the neurons color-coded by clusters in the space spanned by the principal components used for clustering in HDB (left) and VTA (right). **D** Average (colored) and single-unit (gray) waveforms of HDB (top) and VTA (bottom) neurons in each cluster. **E** Average ±SEM spike shape characteristics (valley-to-peak ratio, peak-to-valley time, spike width) and the baseline firing rate distinguish putative cholinergic neurons (teal) from non-cholinergic HDB neurons (left) and putative dopaminergic neurons (purple, and pink) from non-dopaminergic VTA neurons (right). **F** Average normalized PETHs of all HDB (left) and VTA (right) clusters partitioned based on trial outcome (green, hit; red, false alarm; blue, miss; orange, correct rejection). Mustard and grey lines indicate cue and feedback time, respectively.

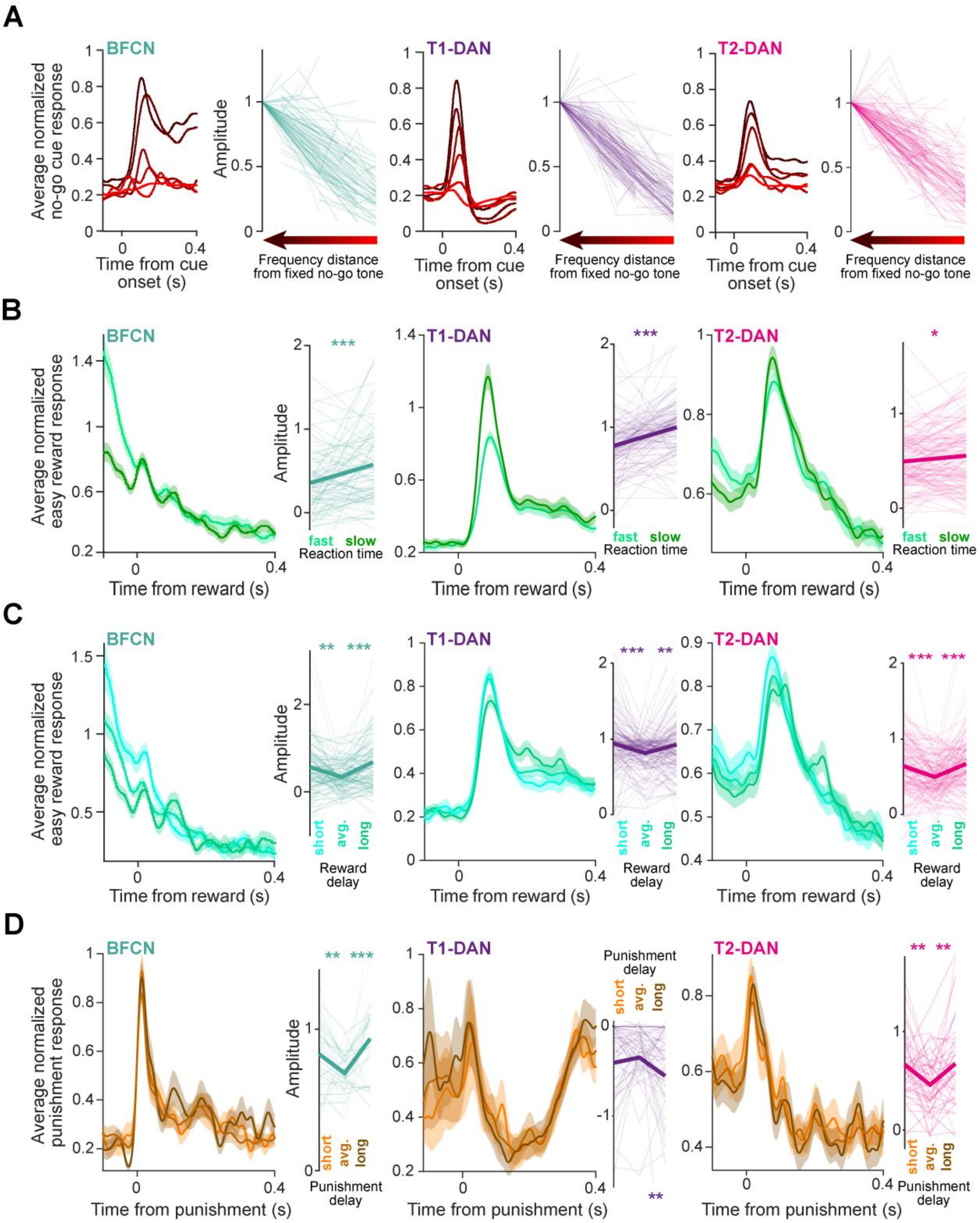


**Figure S6 Cholinergic and dopaminergic neurons show robust signal correlation in various aspects of reward prediction coding.**

**A** Average normalized PETHs of cholinergic (left) and dopaminergic (right) neurons aligned to different no-go cues. Insets, response amplitudes scale with task difficulty indexed by the similarity of the cue frequency to the fixed tones. Lines connect response amplitudes of the same neuron to different tones. **B** Average normalized PETHs aligned to easy rewards revealed larger cholinergic (left) and dopaminergic (right) responses after slow reactions (upper reaction time tercile) compared to fast reactions (lower tercile). Insets, paired comparisons of response amplitudes (BFCNs, p = 0.00087; T1-DANs, p = $9.686 \times 10^{-10}$; T2-DANs, p = 0.0133; Wilcoxon signed rank test). **C** Average normalized PETHs aligned to easy rewards revealed smaller cholinergic (left) and dopaminergic (right) responses after average reward delays. Insets, paired comparisons of response amplitudes (BFCNs, average vs. short delays, p = 0.0036; average vs. long delays, p = $6.217 \times 10^{-6}$; T1-DANs, average vs. short delays, p = 0.00003; average vs. long delays, p = 0.002; T2-DANs, average vs. short delays, p = $2.821 \times 10^{-7}$; average vs. long delays, p = 0.00002; Wilcoxon signed rank test). **D** Same as panel C for hard punishments (BFCNs, average vs. short delays, p = 0.0088; average vs. long delays, p = 0.000997; T1-DANs, average vs. short delays, p = 0.5575; average vs. long delays, p = 0.0054; T2-DANs, average vs. short delays, p = 0.0021; average vs. long delays, p = 0.022; Wilcoxon signed rank test). Error shades represent SEM. *, $p < 0.05$; **, $p < 0.01$; ***, $p < 0.001$.

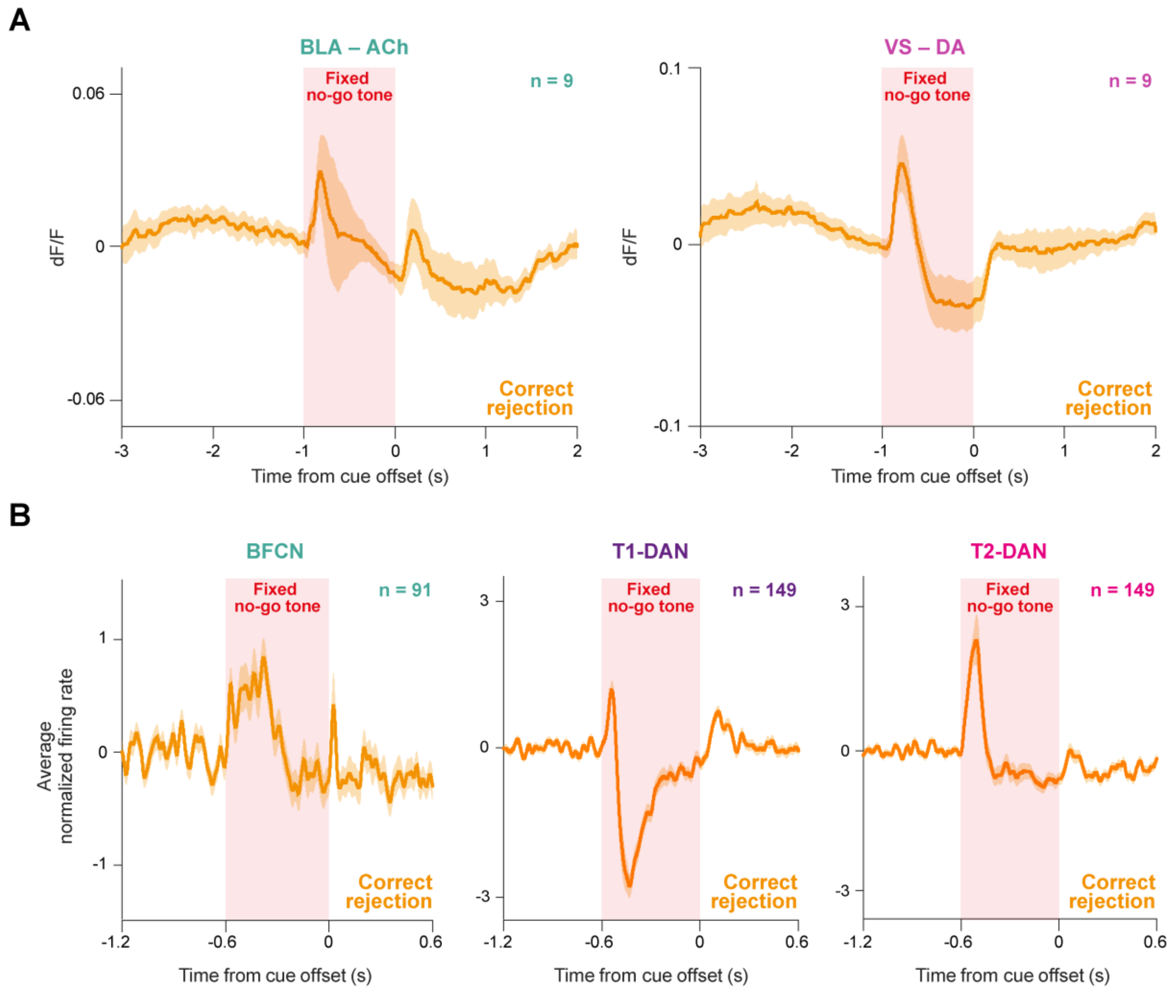


**Figure S7 Avoided punishments lead to an increase in cholinergic and dopaminergic activity.**

**A** Average fluorescent dF/F signals aligned to fixed no-go tone offset in correct rejection trials reveals an increase in both acetylcholine (left) and dopamine (right) release following the successful avoidance of punishments (n = 9 mice). **B** Average Z-scored PETHs aligned to fixed no-go tone offset in correct rejection trials show a moderate increase of BFCN (left), T1-DAN (middle) and T2-DAN firing (right) after avoided punishments. Note that dopamine release and T1-DAN activity is characterized by a brief activation followed by a strong suppression of activity during no-go tones in correct rejection trials, in line with their role in coding signed reward prediction error, which is not apparent in cholinergic and T2-DAN activity. Error shades represent SEM.

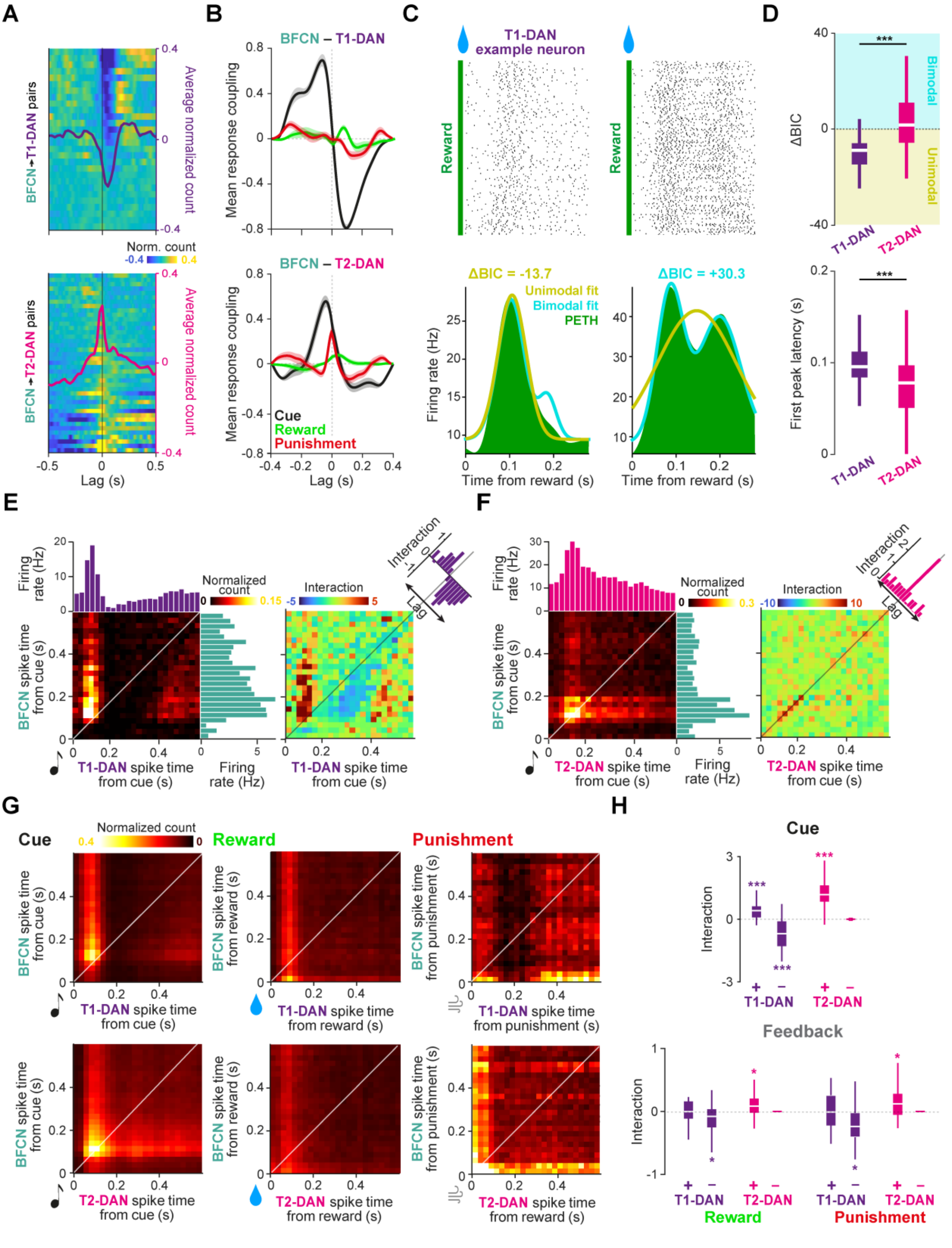


**Figure S8 Cholinergic neurons interact differentially with dopaminergic subtypes.**

**A** Cross-correlograms restricted to spikes not related to behavioral events (inter-trial intervals).Top, cross-correlograms of BFCNs and T1-DANs (rows, pairs of concurrently recorded neurons; blue, low correlation; yellow, high correlation) showing short-latency suppression of T1-DAN activity after BFCN spikes. Bottom, cross-correlograms of BFCNs and T2-DANs showing zero-lag co-activity. Averages are overlaid. **B** Average cross-covariance of normalized PETHs of simultaneously recorded BFCN–T1-DAN (top) and BFCN–T2-DAN (bottom) pairs, aligned to cue (black), reward (green) and punishment (red). Error shades represent SEM. For BFCN–T1-DAN pairs, cue evoked activity is characterized by a robust positive coupling at negative lags (corresponding to faster dopaminergic cue responses preceding BFCN responses), and negative coupling at positive lags, following BFCN activity. Punishment evoked activity is characterized only by the asymmetric negative coupling. Bottom, BFCN–T2-DAN pairs show generally positive coupling temporally modulated by the different response latencies of cues and feedbacks. **C** Reward aligned responses of two example DANs (T1-DAN, left; T2-DAN, right). Top, raster plots revealed that some dopaminergic neurons appeared to exhibit biphasic responses to reward, with two distinct peaks in their activity (see the example neuron on the right). Bottom, corresponding normalized PETHs (green) overlaid with unimodal (single-Gaussian; yellow) and bimodal (two-Gaussian; blue) model fits. The ΔBIC score, defined as the Bayesian information criterion of the unimodal model minus that of the bimodal model, quantifies the relative evidence for the two models; positive values indicate stronger support for the bimodal fit, whereas negative values favor the unimodal fit. **D** Top, T1-DAN activity was predominantly better explained by the unimodal model, whereas bimodal responses were more common among T2-DANs (two-sided Mann-Whitney U-test, $p = 2.78 \times 10^{-21}$). Bottom, distribution of first-response latencies (defined as the earlier peak for bimodal fits and the peak of the single-Gaussian fit for unimodal fits). T2-DANs exhibited faster first responses than T1-DANs (two-sided Mann-Whitney U-test, $p = 7.03 \times 10^{-10}$). Box plots show the median, interquartile range, and non-outlier range. **E** Left, example BFCN–T1-DAN joint PETH (JPETH) with the marginal PETHs shown as bar plots on the sides (right: BFCN; top: DAN). Right, interaction matrix, computed as the JPETH minus the product of the marginal PETHs. Inset, mean interaction along the diagonals of the matrix, reflecting the lag between the two neurons. **F** Same as E for an example BFCN–T2-DAN pair. **G** Average JPETH across all BFCN–T1-DAN (top) and BFCN–T2-DAN (bottom) pairs aligned to cue (left), reward (middle), and punishment (right) reflecting the same temporal dynamics described in panel B. **H** Distribution of interaction strengths around the population average interaction peak (+) and trough (-) lags following cue (left), reward (middle) and punishment (right). Wilcoxon signed-rank tests against zero; $^*p < 0.05$, $^{**}p < 0.01$, $^{***}p < 0.001$. Box-whisker plots show median, interquartile range and non-outlier range.

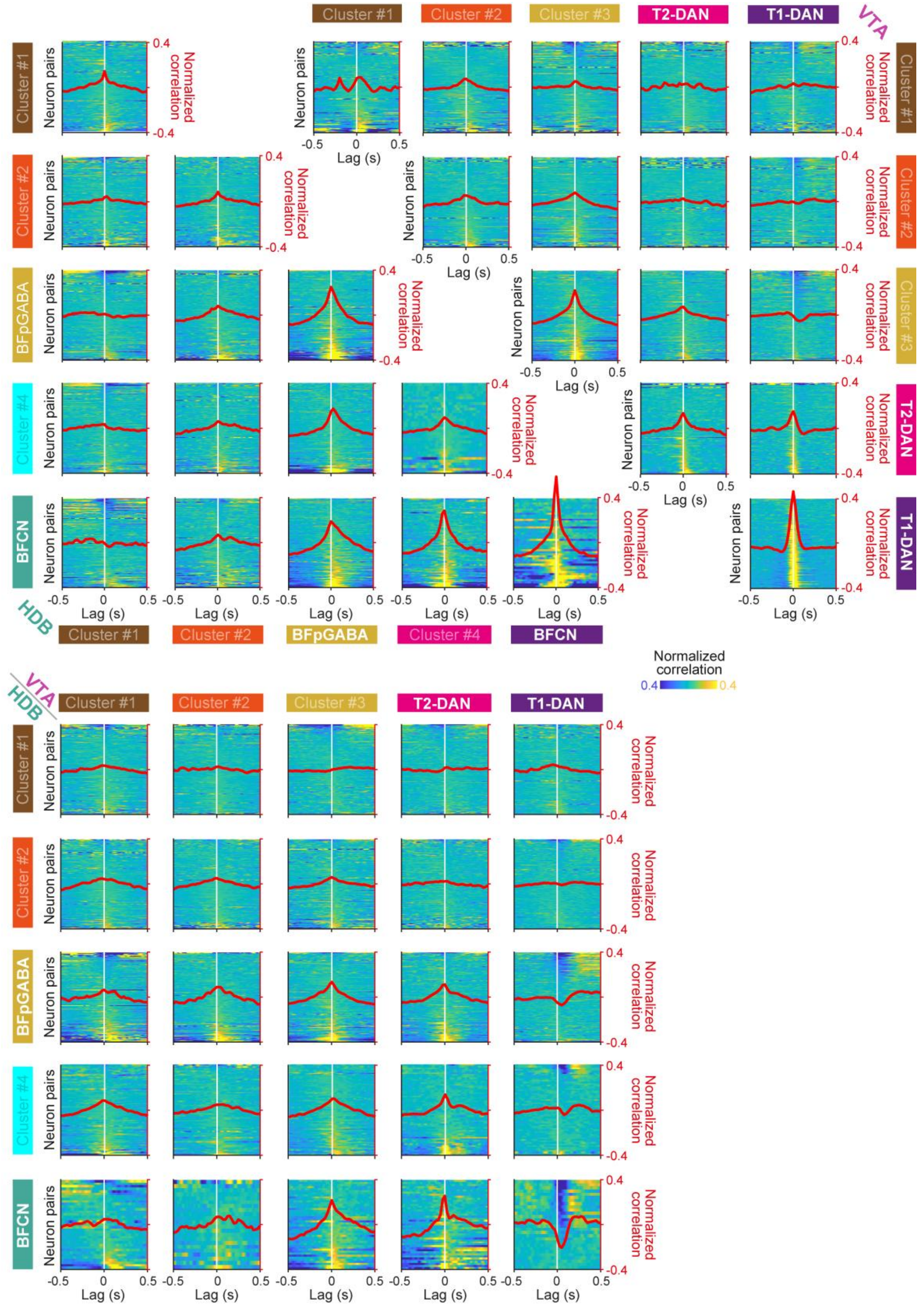


**Figure S9 Cross-correlation structure of neuron clusters in HDB and VTA.**

Cross-correlogram matrix between all neuron clusters (see Fig. S5) within brain areas (top) and across HDB and VTA (bottom). Analyses were restricted to inter-trial intervals to eliminate the potential co-modulatory effects of behavioral events; blue indicates low and yellow indicates high correlation values. Superimposed red graphs represent average correlograms.

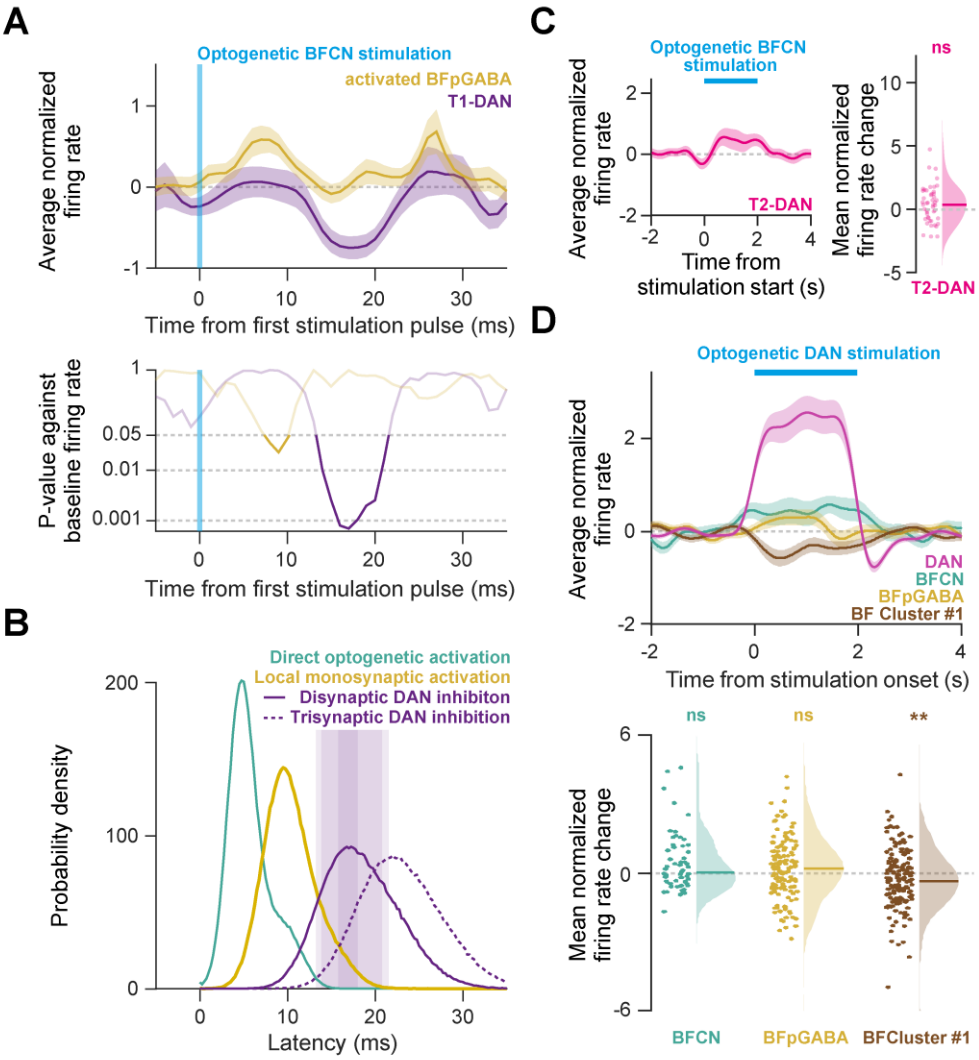


**Figure S10 Optogenetic stimulation experiments support a Type 1-selective disynaptic inhibitory pathway from basal forebrain cholinergic to midbrain dopaminergic neurons.**

**A** Fast effects of optogenetic BFCN stimulation follow a BFCN → BF-GABA → T1-DAN sequence. Top, average normalized peri-event time histograms of T1-DANs (purple) and activated putative basal forebrain GABAergic neurons (BFpGABA, yellow; cluster #3 neurons in Fig. S5A that showed above-baseline firing over the full stimulation train, Fig. 3C). Error shades, SEM. Bottom, p-values of two-sided Wilcoxon signed-rank tests of baseline-normalized firing rate in 1-ms bins as a function of time from the first stimulation pulse (log scale; shaded area: $p > 0.05$, no significant change from baseline). **B** Latency distributions predicted by a Monte Carlo model ($3\times10^5$ samples) for the optogenetic BFCN activation (cyan), the monosynaptic activation of local GABAergic neurons (yellow), a disynaptic BFCN→BF-GABA→T1-DAN pathway (purple) and a trisynaptic pathway with an additional glutamatergic relay (purple dotted). Shaded area: observed T1-DAN reaction latency, shade transparency reflects significance levels of $p < 0.05$, 0.01 and 0.001, see panel A. The disynaptic prediction falls within the observed response window, whereas the trisynaptic prediction was less likely to explain fast measured reactions. BFCN first-spike latency and second-spike probability were measured; the remaining delays were taken from the literature (see Methods). **C** The optogenetic activation of BFCNs had no significant effect on T2-DANs, shown by the average PETH aligned to photostimulation onset (left), and the distribution of firing rate changes (right; $p = 0.1453$, two-sided Wilcoxon signed rank test). **D** Effect of the optogenetic activation of DANs on basal forebrain clusters. Top, average DAN PETH aligned to stimulation onset show that DANs increase their firing upon direct stimulation (tagging). Middle, average basal forebrain PETHs aligned to photostimulation onset. Error shades, SEM. Bottom, no significant firing rate changes were detected in BFCNs ($p = 0.0682$) and putative GABAergic BF neurons ($p = 0.1385$) upon DAN stimulation; however, neurons of the first HDB cluster were significantly inhibited (**, $p = 0.0018$; two-sided Wilcoxon signed-rank test).

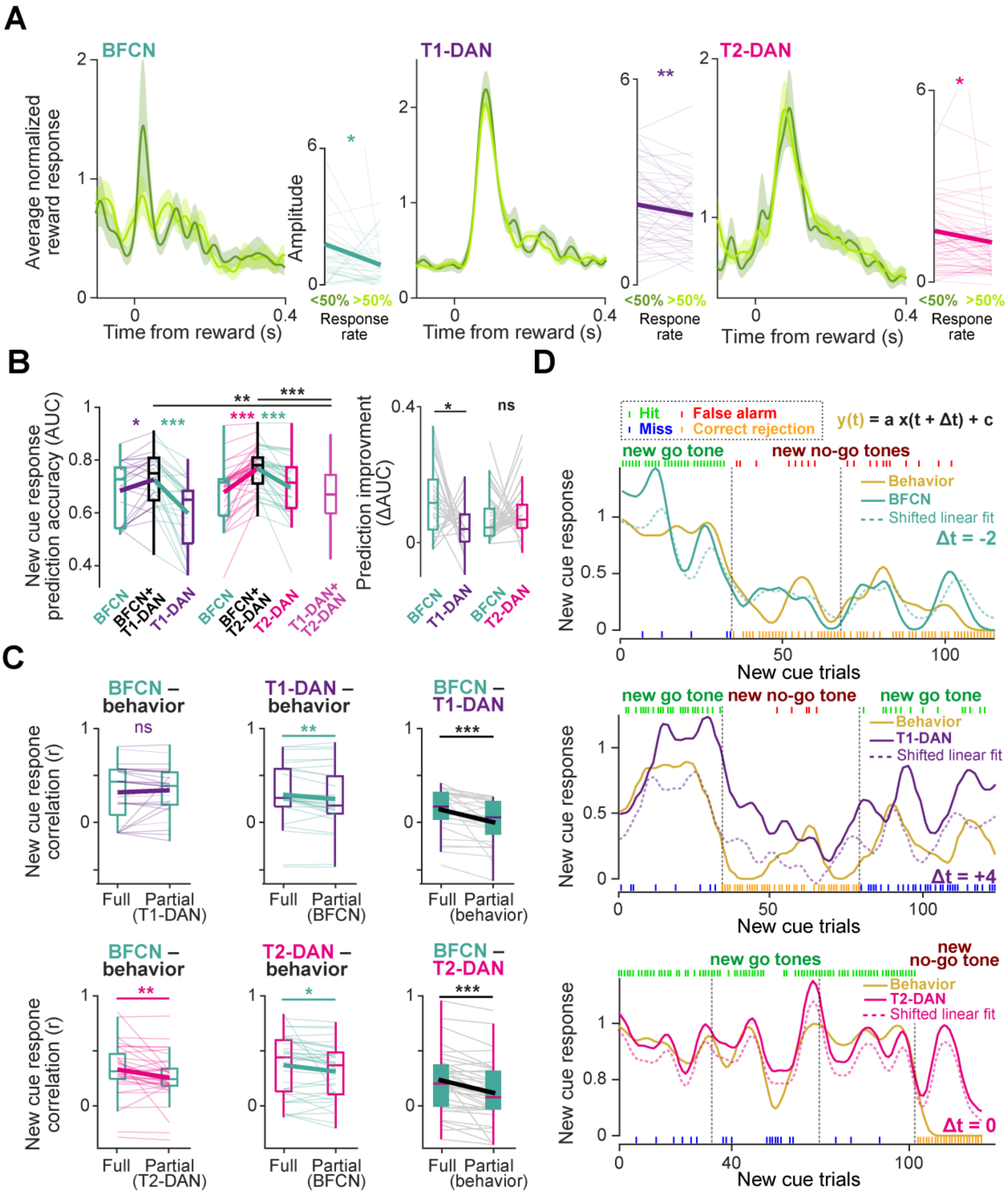


**Figure S11 Cholinergic and dopaminergic neurons jointly predict behavioral decisions, with an asymmetrical contribution from T1-DANs and a symmetrical contribution from T2-DANs.** **A** Average normalized PETHs aligned to hard rewards revealed larger neuronal responses during low (< 50%) responsiveness both in cholinergic (left) and dopaminergic (right) neurons. Error shades, SEM. Inset, paired comparisons of response amplitudes (BFCNs, $p = 0.0374$; T1-DANs, $p = 0.0015$; T2-DANs, $p = 0.0265$; Wilcoxon signed rank test). **B** Left, prediction accuracy of behavioral responses in new-cue trials based on simultaneously recorded cholinergic and dopaminergic single-neuron responses (logistic regression model). BFCNs slightly outperformed Type 1 ($p = 0.023$) but not Type 2 ($p = 0.457$) dopaminergic neurons. Combining the information from different neurons types (BFCN + T1-DAN or BFCN + T2-DAN) outperformed predictions based on single types (BFCN vs. BFCN + T1-DAN, $p = 0.0102$; T1-DAN vs. BFCN + T1-DAN, $p = 4.6 \times 10^{-5}$; BFCN vs. BFCN + T2-DAN, $p = 4.8 \times 10^{-7}$; T2-DAN vs. BFCN + T2-DAN, $p = 4.37 \times 10^{-7}$; two-sided Wilcoxon signed-rank tests) and also predictions based on the combination of the two dopaminergic subtypes (T1-DAN + T2-DAN vs. BFCN + T1-DAN, $p = 0.0088$; T1-DAN + T2-DAN vs. BFCN + T2-DAN, $p = 3.2 \times 10^{-7}$; two-sided Mann-Whitney U-test). Right, BFCNs contributed more to the combined prediction than T1-DANs ($p = 0.023$), whereas for T2-DAN contributions were comparable ($p = 0.457$, two-sided Wilcoxon signed-rank tests). **C** Comparison of full and partial Spearman correlation coefficients (r). Thin lines, single BFCN-DAN pairs; thick lines, average. Left, correlation between BFCN new-cue-responses and behavioral responsiveness, controlled for simultaneously recorded dopaminergic (top, T1-DAN; bottom, T2-DAN) activity. Middle, correlation between dopaminergic (top, T1-DAN; bottom, T2-DAN) responses and behavioral responsiveness, controlled for simultaneous BFCN activity. Right, correlation between BFCN and dopaminergic (top, T1-DAN; bottom, T2-DAN) activity, controlled for behavioral responsiveness. Top row, the correlation between BFCN activity and behavioral responses was not significantly altered by controlling for the variance explained by T1-DANs ($p = 0.374$, Wilcoxon signed-rank tests). In contrast, the correlation between T1-DAN activity and behavior was significantly reduced when controlling for BFCNs ($p = 0.00825$, Wilcoxon signed-rank tests). This asymmetry suggests that BFCNs carry independent behaviorally relevant information while they may exert a modulatory influence on T1-DANs. In line with the latter, correlations between BFCN and T1-DAN activity were significantly reduced after removing behavior-related variance ($p = 5.6 \times 10^{-5}$, Wilcoxon signed-rank tests), often approaching zero or negative values. Bottom row, behavioral correlations of both BFCNs and T2-DANs were significantly reduced (BFCN, $p = 0.00394$; T2-DAN, $p = 0.0147$; Wilcoxon signed-rank tests), but not eliminated, after controlling for the other cell type, indicating partially shared variance. Similarly, the correlations between BFCN and T2-DAN activity were significantly reduced ($p = 6.7 \times 10^{-6}$, Wilcoxon signed-rank tests), but remained positive, when controlling for behavioral responsiveness. **D** Example sessions with BFCN (top, teal), T1-DAN (middle, purple) and T2-DAN (bottom, magenta) responses along with the animal's response rate to novel cues ('behavior', gold; single trial outcomes are also indicated by colored ticks; green, hit trials; red, false alarm trials; blue, miss trials; orange, correct rejection trials). Vertical dashed lines, introduction of new tone-outcome pairings; valence (go vs. no-go) is labelled above each block. Response trends were smoothed by a Gaussian kernel with standard deviation of 1.4 trials; neuronal responses were normalized to fixed go tone responses. Colored dashed traces, a lagged linear regression model $y(t) = a\,x(t + \Delta t) + c$ was fitted to explain behavioral responsiveness trends $y(t)$ from time-shifted ($\Delta t$) neuronal activity $x(t)$ (dashed colored lines). $\Delta t$ values indicate the time-shift of the best model fit (negative $\Delta t$, neuronal activity anticipated behavioral trends; positive $\Delta t$, neuronal activity lagged behavioral trends). Box-whisker plots show median, interquartile range and non-outlier range. $^{*}p < 0.05$, $^{**}p < 0.01$, $^{***}p < 0.001$.

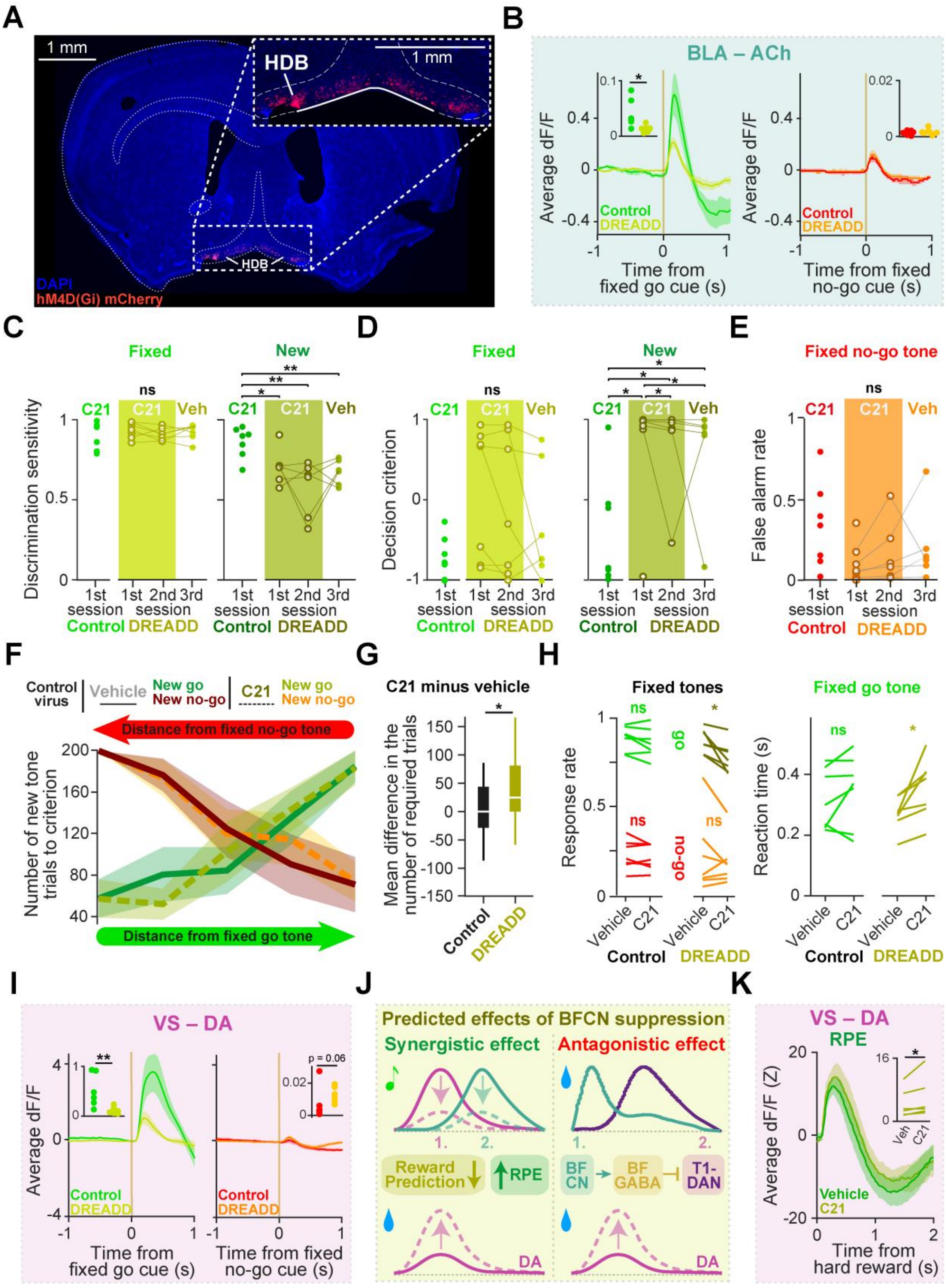


**Figure S12 Chemogenetic inhibition of basal forebrain cholinergic neurons.**

**A** Fluorescent micrograph of the bilateral HDB injection sites. Red, mCherry in neurons expressing hM4D(Gi) DREADD; blue, DAPI.

**B** Average ACh release in the BLA aligned to the fixed go (left) and no-go (right) cue in control (green and red, respectively) vs. DREADD-expressing (yellow and orange, respectively) animals (first session with novel association; error shades, SEM). Inset, fixed go cue-evoked ACh release was reduced in DREADD-expressing mice (p = 0.026; two-sided Mann-Whitney U-test). **C** Nonparametric discrimination sensitivity of the fixed (left) and the new (right) go tone versus the fixed no-go tone. DREADD-expressing mice receiving the C21 DREADD ligand maintained good discrimination of the previously learned fixed go tone (compared to control-virus injected mice: 1st new cue session, p = 0.3829; 2nd new cue session, p = 0.535; 3rd new cue session with vehicle injection, p = 0.4452; two-sided Mann-Whitney U-test) but impaired discrimination learning of the novel go tone (compared to control-virus injected mice: 1st new cue session, p = 0.0379; 2nd new cue session, p = 0.0047; two-sided Mann-Whitney U-test). On a control day with vehicle injection (following two C21 sessions), the deficit still persists compared to control-virus injected mice (p = 0.0047, two-sided Mann-Whitney U-test). **D** Same as C for nonparametric response bias. DREADD-expressing mice receiving the C21 DREADD ligand showed no difference in response bias for the fixed go tone (compared to control-virus injected mice: 1st new cue session, p = 0.1282; 2nd new cue session, p = 0.3829; 3rd new cue session with vehicle injection, p = 0.4452; two-sided Mann-Whitney U-test), but adopted a more conservative bias for the novel go tone (1st new cue session, p = 0.0262; 2nd new cue session, p = 0.0221; 3rd new cue session with vehicle injection, p = 0.0221; two-sided Mann-Whitney U-test). **E** DREADD-expressing mice receiving the C21 DREADD ligand maintained good performance on the previously learned fixed no-go association (compared to control-virus injected mice: 1$^{st}$ new-cue session, p = 0.0728; 2$^{nd}$ new-cue session, p = 0.2086; 3$^{rd}$ new-cue session with vehicle injection, p = 0.366; two-sided Mann-Whitney U-tests). **F** In mice injected with control virus, psychometric learning curves (see Fig 1.b) in sessions with C21 vs. vehicle injections revealed no learning differences in control mice. Lines, grand-averages; error shades, SEM. **G** Learning deficits in session with C21 injections were significantly larger in DREADD-expressing than in control animals (p = 0.0215, Mann-Whitney U-test). Distribution of the intermediate-difficulty task (i.e., tones >1 kHz away from the fixed tones) is plotted. Box-whisker plots show median, interquartile range and non-outlier range. **H** Hit rate after fixed go-tones and reaction time were slightly altered in sessions with C21 injection in the DREADD-expressing (hit rate, p = 0.0156; reaction time, p = 0.0156), but not in control mice (hit rate, p = 0.2188; reaction time, p = 0.4688). False alarm rate after fixed no-go tones did not change significantly (DREADD, p = 0.6875; control, p = 0.6875; Wilcoxon signed rank test). **I** Same as panel B for DA release in the VS. Go cue-evoked release was reduced (p = 0.0087), whereas no-go cue-evoked release showed a trend toward an increase (p = 0.0649) in DREADD-expressing mice (two-sided Mann-Whitney U-test). **J** Expected effect of cholinergic suppression on dopaminergic reward responses. Left, suppressing BFCNs weakens synergistic dopaminergic reward predictions, yielding higher reward prediction error in rewarded trials, expected to be reflected in increased DA release following rewards. Right, rapid activation of BFCNs following reward precedes T1-DAN responses, allowing disynaptic inhibition of T1-DANs (Fig. 3); suppressing BFCNs should therefore release T1-DANs from this inhibition, likewise increasing dopamine release after rewards. **K** Average DA release aligned to hard rewards from C21 vs. vehicle sessions of DREADD-expressing mice. Error shade, SEM. Inset, rewards were followed by enhanced DA release in C21 sessions (p = 0.0312, two-sided Wilcoxon signed-rank test). *, p < 0.05; **, p < 0.01; ***, p < 0.001.
